# Structure of RADX and mechanism for regulation of RAD51 nucleofilaments

**DOI:** 10.1101/2023.09.19.558089

**Authors:** Swati Balakrishnan, Madison Adolph, Miaw-Sheue Tsai, Kaitlyn Gallagher, David Cortez, Walter J. Chazin

## Abstract

Replication fork reversal is a fundamental process required for resolution of encounters with DNA damage. A key step in the stabilization and eventual resolution of reversed forks is formation of RAD51 nucleoprotein filaments on exposed ssDNA. To avoid genome instability, RAD51 filaments are tightly controlled by a variety of positive and negative regulators. RADX is a recently discovered negative regulator that binds tightly to ssDNA, directly interacts with RAD51, and regulates replication fork reversal and stabilization in a context-dependent manner. Here we present a structure-based investigation of RADX’s mechanism of action. Mass photometry experiments showed that RADX forms multiple oligomeric states in a concentration dependent manner, with a predominance of trimers in the presence of ssDNA. The structure of RADX, which has no structurally characterized orthologs, was determined *ab initio* by cryo-electron microscopy (EM) from maps in the 2-3 Å range. The structure reveals the molecular basis for RADX oligomerization and binding of ssDNA binding. The binding of RADX to RAD51 filaments was imaged by negative stain EM, which showed a RADX oligomer at the end of filaments. Based on these results, we propose a model in which RADX functions by capping and restricting the growing end of RAD51 filaments.

**Significance:** Despite the central role of RAD51 in DNA replication and repair processes, the mechanisms of action of its many modulators are poorly understood. Here we combine structural and biophysical data to determine how the negative regulator RADX functions. We show that RADX oligomerizes upon binding DNA, and caps RAD51 filaments at the ends to prevent extension. This work advances knowledge of how RAD51 filaments can be modulated to regulate replication fork reversal and maintain genomic stability.

## Introduction

Replication fork reversal is a crucial aspect of genome maintenance, allowing for either replication coupled repair or damage bypass upon encounter of a DNA lesion^1^. Fork reversal is tightly controlled, with multiple proteins regulating both the formation and stabilization of reversed forks. Unregulated fork reversal in humans is seen to slow down the process of replication, increase the incidence of double-strand breaks (DSBs) and lead to nascent strand degradation, all of which can result in genome instability^2^.

The single-strand DNA binding (SSB) recombinase RAD51 has a central role in this process, promoting fork reversal^3^ and preventing nascent strand degradation at persistently stalled forks^4,5^. It may also promote strand invasion to restart stalled forks^6^. Upon replication fork stalling, the exposed single strand DNA (ssDNA) is protected by RPA, the ‘first-responder’ SSB. RAD51 must then replace RPA, a process driven by mediators such as BRCA2 that load and stabilize RAD51 nucleoprotein filaments^7^. Inappropriate replication fork reversal can be deleterious to genome stability, so RAD51 action is tightly regulated by multiple proteins. BRCA2 and RAD51 paralog proteins are viewed as positive regulators of formation of RAD51 filaments on ssDNA, even though the exact mechanism of action for the RAD51 paralogs remains unknown^8,9^. RADX (*R*PA-related R*AD*51-antagonist on *X*-chromosome) was recently identified as a modulator of RAD51 at replication forks^10^. Its functional relevance is underscored by the observation that RADX expression levels are inversely correlated to the PARP-inhibitor resistance of BRCA2-deficient cells, and that higher levels of RADX indicate better outcomes in breast and lung cancer patients^10^. Interestingly, patients with high levels of RAD51 fare poorly when afflicted with the same cancer types, suggesting that the modulation of RAD51 function by RADX can impact responses to treatments directed towards DNA replication and repair. This also makes RADX a potential target for the development of future therapeutics.

Previous investigations into RADX function show that RADX allows cells to maintain a high capacity for homologous recombination, while buffering RAD51 between fork reversal and protection to ensure replication integrity^10–13^. Additionally, pull-down and electrophoretic mobility shift assays have identified that RADX binds to DNA and RAD51, and mutants have been designed based on sequence homology, structural modeling and biochemical approaches to parse the importance of these interactions for RADX function^10,12^. Two models for the mechanism of action of RADX have been proposed as a result of these studies. Single molecule fluorescence experiments led to a model proposing sequestration of DNA by RADX as a mechanism of action^14^. The second model is based on our biochemical analysis, which showed that the ATP hydrolysis rate of RAD51 increases in the presence of RADX^12^. Since hydrolysis of ATP by RAD51 leads to release from DNA, this observation implies that RADX functions by promoting filament disassembly.

In order to better understand how RADX participates in fork remodeling and in particular, modulates formation and disassembly of RAD51 filaments, we set out to determine the structural and molecular mechanisms of RADX function. It has been proposed that RADX oligomerization may be crucial to its function^15^; we used mass photometry to test this hypothesis and systematically characterize the concentration and ssDNA dependence of the population distribution of oligomeric states of RADX. We went on to determine high resolution cryo-electron microscopy (cryo-EM) structures of the predominant RADX trimer and of the secondary population of RADX tetramer, both in the presence of ssDNA. We also used EM to investigate the binding of RADX to RAD51 filaments. Together, these results, integrated with all previous data, provided the basis for proposing that RADX function in fork remodeling involves RADX oligomers binding to and capping the ends of RAD51 filaments.

## Results

### RADX oligomerization is stabilized and preferentially forms trimers upon binding ssDNA

RADX forms homo-oligomers and this property is required for full function^14,15^. RADX mutants that cannot properly oligomerize were seen to cause signs of replication stress in cells, despite retaining the ability to bind ssDNA and localize to replication forks at levels comparable to wild-type RADX. The experiments performed previously demonstrated that RADX forms oligomers but did not specifically characterize which oligomeric states are present.

Mass photometry is a powerful method to characterize the mass of particles present in a solution and was applied to quantify RADX oligomerization under different conditions. RADX alone was found to be primarily monomeric at a concentration of 50 nM, but a modest two-fold increase in concentration to 100 nM resulted in a reduction of monomer in favor of the dimer and larger oligomeric states (Fig. 1a). Importantly, the higher order oligomeric states are not discrete well-defined states, but rather appear as an amorphous histogram (Fig. S1a). This is indicative of rapid transient association/dissociation on the timescale of the mass photometry measurement, where no specific oligomer is stable enough to be cleanly isolated. Thus, RADX alone exists in a concentration-dependent equilibrium between a wide range of oligomeric states.

**Figure 1.**
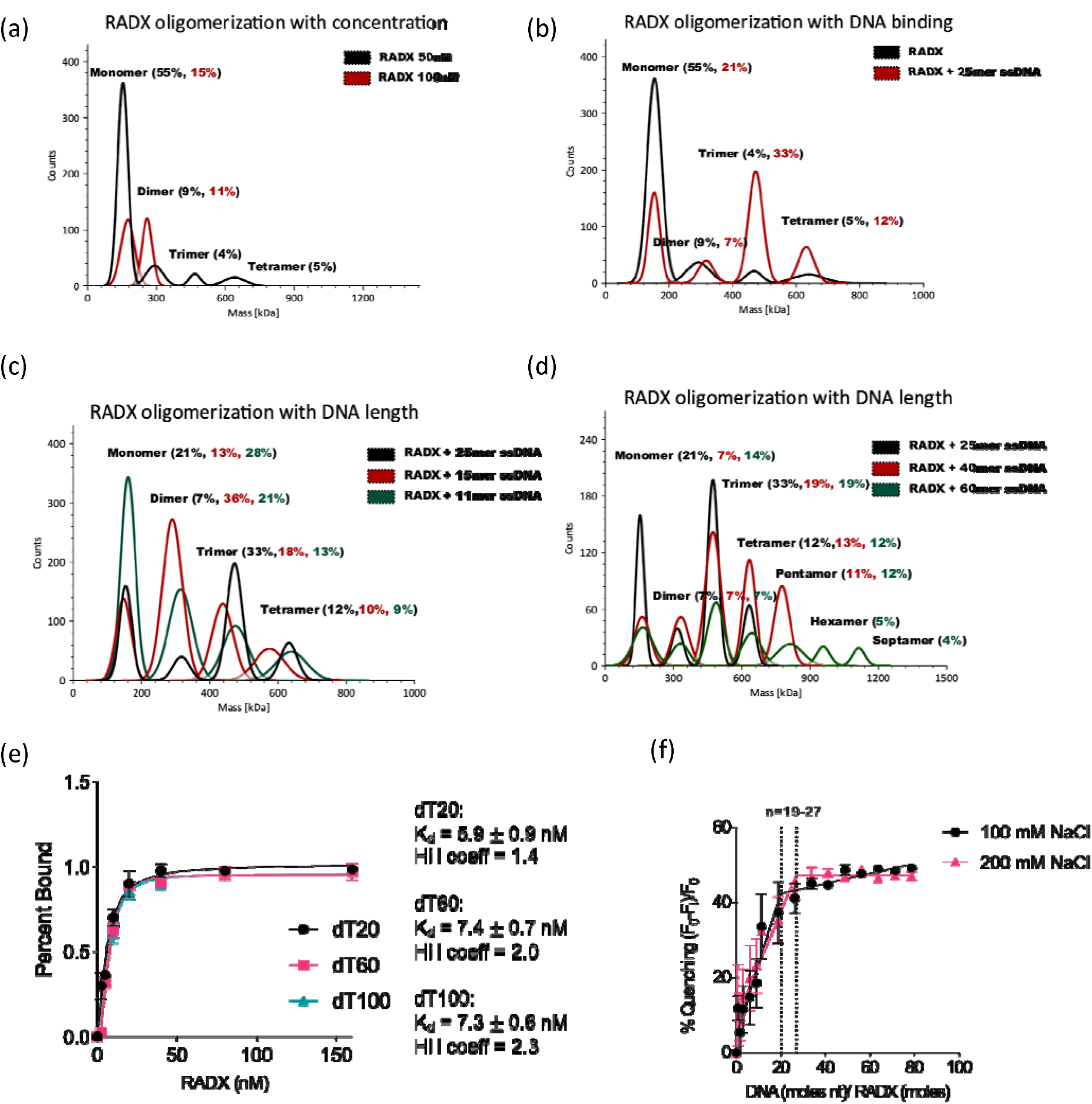
RADX oligomerization is stabilized and preferentially forms trimers upon binding ssDNA. (a) RADX is primarily monomeric at 50 nM but dimerizes at 100 nM. (b) Addition of ssDNA to monomeric RADX leads to a significant increase in the population of trimers, from 4% to 33%. (c),(d) Addition of different lengths of ssDNA leads to varying distribution of oligomers. (e) DNA binding affinity of RADX determined by fluorescence polarization anisotropy show there is no dependence on the length of the substrate. (f) Tryptophan quenching assay measuring the DNA footprint of RADX shows a footprint length of 19-27 nucleotides, consistent with the mass photometry results in panel (c).

To elucidate the effect of DNA binding on RADX oligomerization, mass photometry data was acquired for 50 nM RADX in the presence of a variety of ssDNA oligomers. Binding of dT25 had a significant effect on the oligomerization of RADX, producing a prominent trimer along with discrete dimers and tetramers (Fig. 1b). To ensure the results were not unique to this polypyrimidine oligomer, additional experiments for mixed sequence 25-mer ssDNA were acquired. The data obtained for all 25-mers were strikingly similar (Fig. S1.b), which shows that RADX binds ssDNA with no sequence specificity. Thus, although RADX exists as a large and heterogeneous equilibrium distribution of oligomerization states in the absence of ssDNA, the addition of ssDNA significantly stabilizes specific oligomerization states.

In order to determine if this observation was dependent on the length of the substrate, additional mass photometry data were acquired for 50 nM RADX bound to varying lengths of ssDNA (Fig. 1). The data show that the distribution of oligomeric states is dependent on the length of DNA available for binding (Fig. 1). RADX primarily forms trimers when bound to dT25 and dT20, but the trimers are apparently destabilized as the ssDNA is shortened as dimers become the dominant states in the presence of dT15 (Fig. 1c) and dT11 shifts the equilibrium further towards the monomer. The presence of longer ssDNA lengths such as dT40 and dT60 lead to the formation of larger RADX-DNA complexes with more (e.g., four, five, six) RADX’s bound to the ssDNA (Fig. 1d). A striking observation in these experiments was that even in the presence of these longer lengths of ssDNA the trimer remains the most abundant state. These results show that the trimer is more stable than other oligomeric states and indicate the higher mass states are not higher order oligomers but rather two or more lower order oligomers (e.g., 2X trimers) loaded on the ssDNA.

To further ensure that any effects observed were not due to differences in the mode of binding ssDNA, we measured the affinity of RADX using a fluorescence anisotropy assay with fluorescein labeled dT20, dT60, and dT100 (Fig. 1e). The K_d_ values measured were all very similar (5.9 ±0.9 – 7.4 ±0.6 nM) indicating a consistent ssDNA binding mode. Additionally, the value of the Hill coefficient was seen to be above 1 even for dT20, indicating that energetic coupling leads to cooperativity in RADX oligomerization and binding of ssDNA. This binding model is considerably more complex than the one involving only monomeric RADX, and indicates that the K_d_ determined here is an apparent K_d_. We were also able to establish that the DNA footprint is 19-27 nucleotides by monitoring the fluorescence quenching of tryptophan residues in RADX binding to poly-dT, which is consistent with the mass photometry data showing that the trimer and tetramer are the dominant states when bound to ssDNA.

Mass photometry, while powerful, is restricted to measurements in the nM range of concentrations. To further explore RADX oligomerization at higher concentrations, we performed small angle X-ray scattering (SAXS) experiments. It is not possible to concentrate RADX above 5 μM in the absence of ssDNA and as a result we were unable to collect high quality data to perform SAXS experiments on the free protein. However, RADX is more readily concentrated in the presence of ssDNA and has a much lower tendency to aggregate, so SAXS experiments could be performed for the complex with dT25. Linearity in the Guinier region and the ability to include data points to very low values of q (n_min_=2) indicated the data were of high quality. Rg parameters extracted directly from the data using the Guinier fit (q_min_Rg = 0.79, q_max_Rg = 1.35) also indicated the data are of high quality and that the protein is globular in nature. The calculation of the mass of the complex of RADX with dT25 at this concentration from the SAXS data corresponds to a tetrameric state (Fig. S1c). The 50-fold increase in concentration from 100 nm to 5 μM appears to cause a shift from a preference for trimer to tetramer, which is consistent with the trend of higher order oligomers with increasing protein concentration. We note that the molecular envelope calculated from the SAXS data fits reasonably well to the high-resolution cryo-EM structure of the tetramer that is described below (Fig. S1c).

### High resolution structure shows RADX is comprised of four independent OB-fold domains

While significant data has been accumulated about RADX function, the lack of a 3D structure has limited the ability to draw firm conclusions about its mechanism of action. A previous analysis of the RADX sequence proposed that RADX contains three N-terminal OB-fold domains and a dual motif C-terminal domain^10^. However, the lack of RADX orthologs and its poor sequence homology to other proteins with known structures precludes reliable prediction of its 3D structure, although we note that AlphaFold does produce a structural model for monomeric RADX with relatively high confidence for 75% of the protein. We therefore set out to determine the structure of RADX by cryo-EM.

Initial screening was performed by selecting specific fractions of the complex of RADX with dT25 purified by size exclusion chromatography. Negative stain imaging of these fractions produced 2D class averages with different sizes and shapes of particles (Fig. 2a). Upon screening the same sample in cryo-EM, the distribution of individual particles was more readily determined, but the resulting 2D class averages remained poorly resolved despite extensive attempts to optimize the data analysis parameters (Fig. 2b). From these results, we surmised that RADX has significant inter-domain flexibility under these conditions.

**Figure 2.**
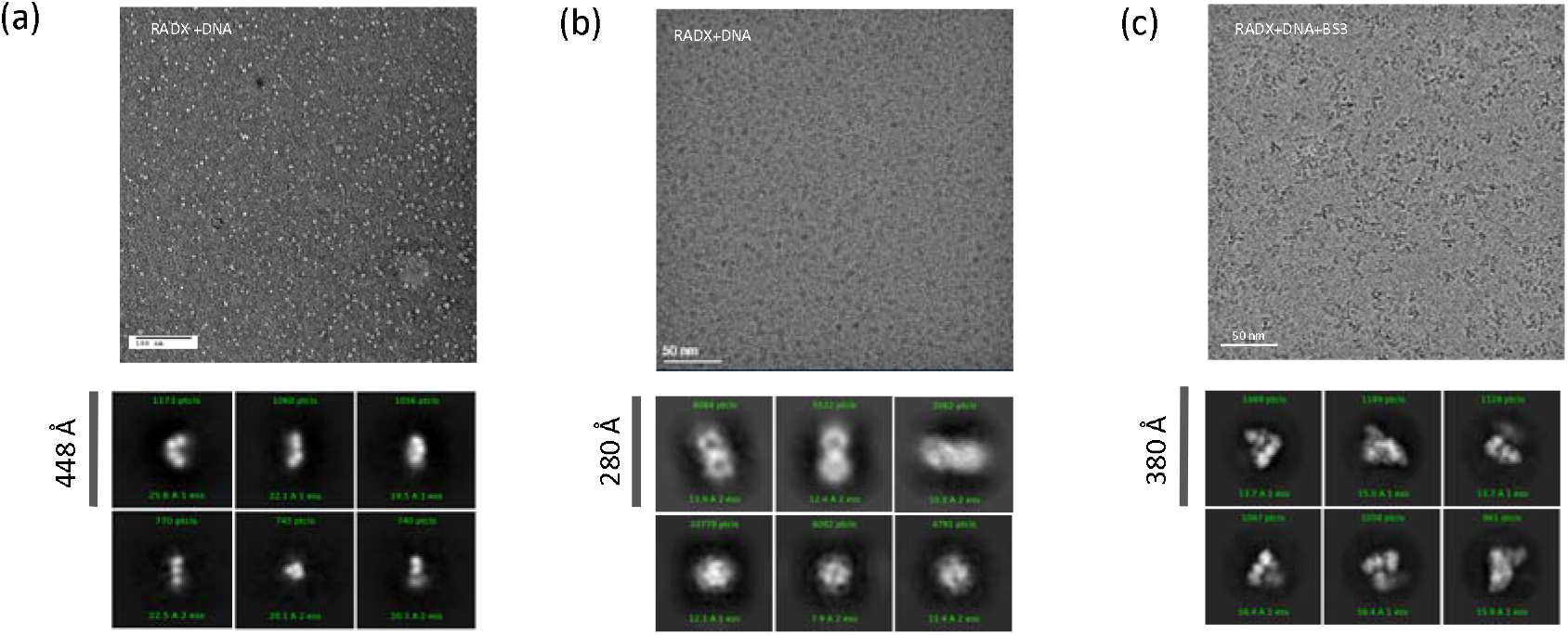
Optimization of RADX for cryo-EM. (a) Negative stain EM micrograph image of the RADX-dT25 complex. 2D classes showing monomeric RADX in a variety of configurations is shown below the micrograph, indicating heterogeneity remains an issue. (b) Cryo -EM micrographs of RADX-dT25 complex without cross-linking. 2D classes under the micrographs show that without cross-linking RADX has very poorly resolved 2D classes due to its flexibility. (c) Cryo-EM micrograph of RADX-dT25 complex crosslinked with BS3. 2D class averages show the resolution improves significantly upon cross-linking, allowing for 3D reconstruction.

To overcome this barrier, samples of the RADX-dT25 complex were chemically cross-linked with BS3, which resulted in significant improvement in the resolution of the 2D classes (Fig. 2c). Particles in these grids were still heterogeneous, which was attributed to population of different oligomeric states. After extensive 2D classification, it became clear that trimer and tetramer particles were dominant and in sufficient quantity to warrant pursuing refinement of their respective structures (Fig. S2, Table S1). Trimer and tetramer particles were then separated out based on both 2D and 3D classification. The distribution of particles from this analysis was ∼70% trimer and ∼30% tetramer, consistent with the ratio observed in the mass photometry data on the non-crosslinked protein (Fig. 1a). This suggests that the trimer and tetramer particles are not artefacts of cross-linking.

The structures of the trimer and tetramer were each determined using a combination of non-uniform refinement and local refinement with masking (Fig. S2). Local refinement was used to circumvent the loss of resolution due to motion between each RADX protomer relative to the others in the oligomer. Thus, separate local refinements were performed on the A-B and B-C RADX pairs within the trimer (Fig. 3a). The maps were then merged to generate the final composite map, which has an average resolution of 2.9 Å (Fig. 3a). The data for the tetramer were processed in the same manner with separate local refinements performed on the A-B, B-C and C-D pairs. The density for the fourth RADX protomer (D) was relatively poor compared to A, B, and C and when combined with the smaller number of particles, resulted in the average resolution of the final map for the tetramer being 3.7 Å (Fig. 3b). Importantly, density for the ssDNA is well resolved in both maps. Since the resolution of the final map was higher, subsequent analyses are based on the structure of the trimer.

**Figure 3.**
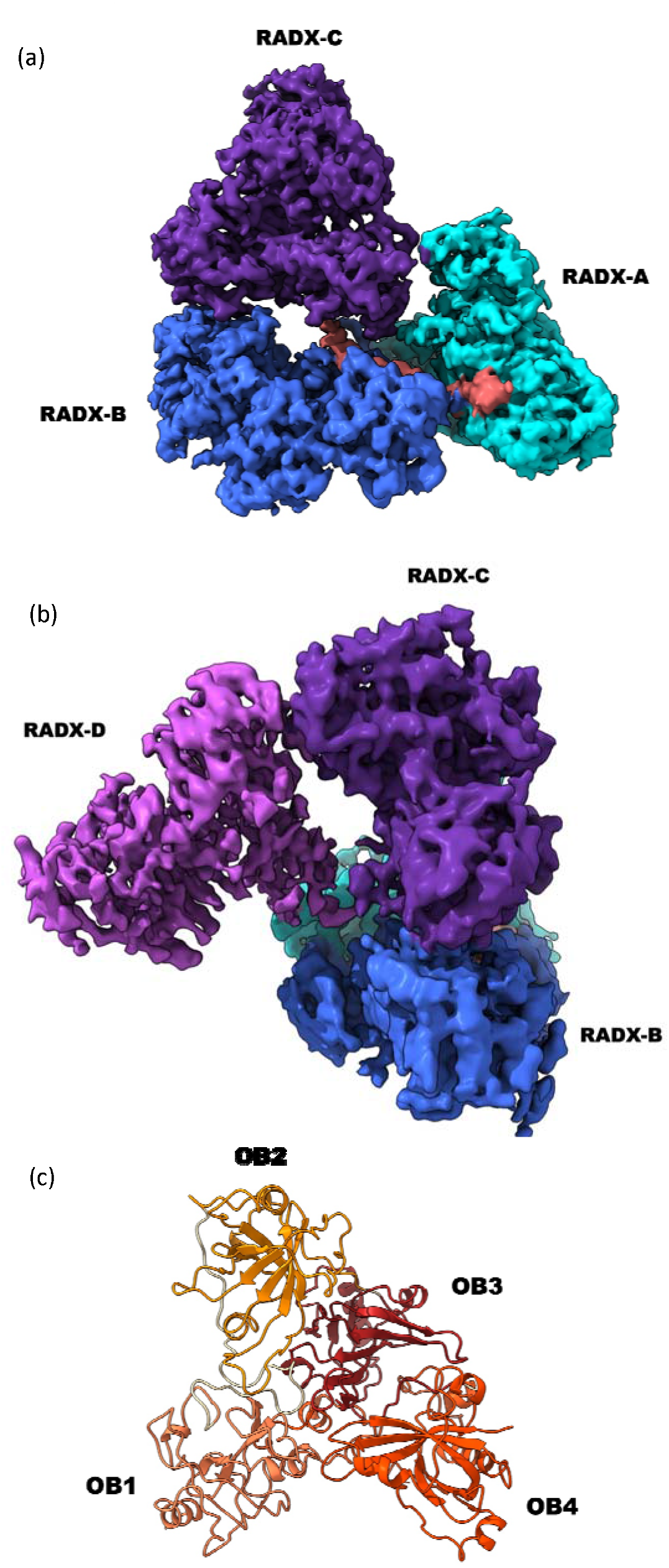
High resolution structure shows RADX is comprised of four independent OB-fold domains. (a) Structure of the RADX trimer bound to ssDNA (pink) with the three protomers in cyan, blue and purple. (b) The structure of the RADX tetramer with the fourth RADX protomer RADX-D (violet). (c) Ribbon diagram of a RADX protomer showing the location of each of the four OB-fold domains.

RADX is found to consist of four OB-fold domains (OB1-OB4) (Fig. 3c), with a 110-residue disordered insertion within the fourth domain spanning S566–I676, for which no density was observed. All four domains are canonical OB-folds with a core β-sheet organized into two three-stranded anti-parallel sub-sheets that are each composed of a central curved β-strand with anti-parallel strands on either side. Each domain also includes 2 or 3 α-helices, with the first helix packed at the bottom of the β-sheet, completing the canonical OB-fold domain structure^42^. There is clear density for the ssDNA in the RADX trimer, which contacts 16 of the 25 nucleotides. Notably, the density of the fourth protomer (RADX-D) in the tetramer is insufficient to visualize all the ssDNA. A key feature of both the trimer and tetramer structures is the lack of symmetry with respect to the three or four protomers within in each structure.

A comparison of RADX to known structures using the programs DALI^40^ and FoldSeek^41^ show that RADX has a unique protein fold, with no alignment matches for all four domains. Many matches are seen for the single domains, such as the alignment of RADX OB1 with RPA 70N (Fig. S3) or OB3 with POT1. However, OB4 stands out as it does not have any well-matched homologs. All potential alignments to OB4 show very large RMSDs (over 10 Å) and very low Z-scores (below 3). The unique combination of domains as reflected in the lack of well-matched homologs make our RADX structure a valuable addition to the evolving understanding of protein structure.

### RADX oligomerization is stabilized by multiple inter-domain interfaces

Oligomerization is a fundamental biochemical property of RADX, so elucidating the driving force for RADX oligomerization is important for understanding its function. To this end, we analyzed the interfaces between RADX protomers in the high-resolution structure of the trimer using the PDBePISA server^43^. The primary interface mediating oligomerization is between OB1 and OB4, with OB4 of RADX-A interacting with OB1 of RADX-B, OB4 of RADX-B interacting with OB1 of RADX-C, and so on. Although the interfaces are the same, differences in the relative orientation of the protomers are required so that each can align successively for oligomerization (Fig. 4a). The three OB1-OB4 interfaces in the trimer have on average a buried surface area of ∼600 Å^2^ and 6 hydrogen bonds or salt bridges (Fig. 4a). Key hydrogen-bonding residues include E526, L529, Q553, N759 and E761 from OB4, and R58, Y138, E140, K141 and R142 from OB1.

**Figure 4.**
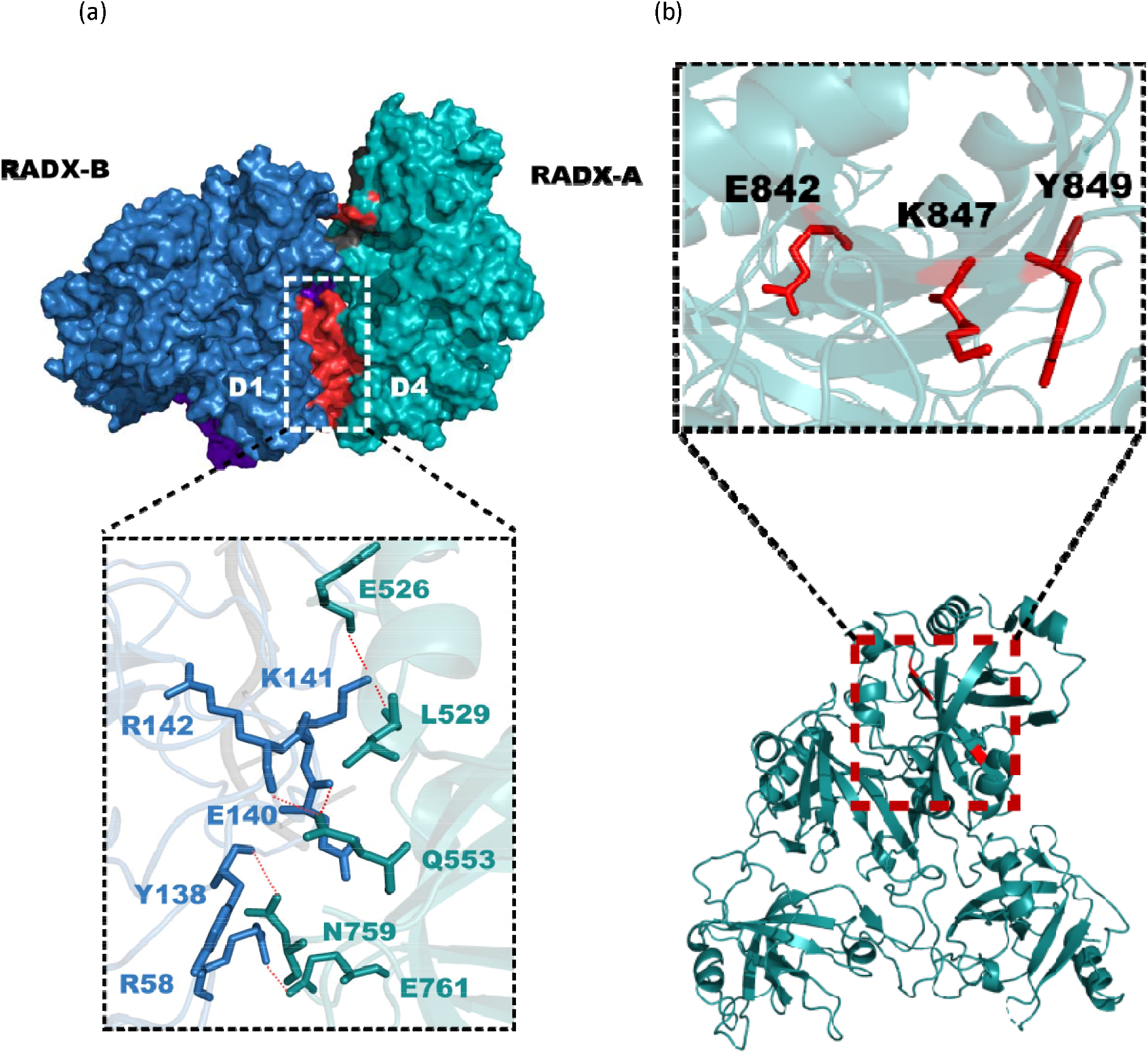
RADX oligomerization is stabilized by multiple inter-domain interfaces. (a) The D1-D4 interaction between two RADX protomers A and B (teal and blue, respectively). RADX oligomerizes primarily via hydrogen bond and salt-bridge interactions between domains 1 and 4. The inset shows the residues involved in hydrogen bonding colored according to the protomer of origin, with the hydrogen bonds shown as red dashed lines. (b) Three of the sites of RADX oligomerization mutations in the central beta sheet of OB4 displayed in red in the inset. Mutation of these residues are likely to perturb the beta sheet and the entire D4 domain, which could contribute to effects observed in functional assays.

A secondary oligomerization interface is provided by successive OB2 domains. These OB2-OB2 interactions are smaller, averaging only ∼100 Å^2^ in buried surface area and with only 1 or 2 hydrogen bonds involving residues R232 and Y307. Remarkably, this secondary interface appears to be sufficient to mediate some degree of oligomerization of RADX as mass photometry data obtained for an N-terminal construct spanning R43-K560 (RADX-N) that lacks the OB4 domain show this truncated protein still forms primarily dimers at a concentration of 50 nM (Fig. S4a). Notably, this RADX-N construct is unable to form any higher order oligomers even in the presence of ssDNA (Fig. S4b), indicating that OB4 is required for their formation.

A previous investigation of RADX function also surmised that the fourth predicted domain in the C-terminus was important for oligomerization, and three single site mutations (E842K, K847E, Y848A) were selected for investigation^15^. These variants inhibited oligomerization, retained DNA and RAD51 binding, and localized to replication forks. Remarkably, the structure now shows that these three residues are not located at the OB1-OB4 oligomerization interface. Rather, all three form part of a β-strand that is critical to the central β-sheet of OB4 (Fig. 4b), which suggests the mutations may disrupt the structural stability of OB4. Future analysis of structure-based variants that specifically disrupt oligomerization will provide confirmation of our earlier studies and more in-depth analysis of the functional role(s) of RADX oligomerization.

### RADX binding of ssDNA is coupled to oligomerization

The discovery and initial analyses of RADX revealed homology to the large subunit of replication protein A (RPA70) and tight binding of ssDNA^10^. RPA binds ssDNA using multiple OB-fold domains^44^, including three in RPA70 (RPA70ABC). The initial prediction of 3 OB-fold domains in RADX suggested that RADX uses the same multi-valent mode of binding as RPA. Subsequent mutational analysis revealed the central role of OB2^10^, but exactly how RADX binds ssDNA remained poorly understood.

The high-resolution structure of RADX shows that it does bind ssDNA in a multi-valent mode, but not in same manner as RPA. Instead, RADX binds ssDNA almost exclusively through OB2 and uses coupling to oligomerization to generate multi-valency and attain high affinity (Fig. 5a). In the RADX trimer, all three protomers are aligned in a manner that allows for tight binding of the ssDNA via their respective OB2 domains, with direct contacts to 16 nucleotides. This observation is consistent with the 19-27 DNA footprint obtained for poly-dT, given that excluded site size determined by footprinting is usually larger than the number of residues in direct contact and that the RADX tetramer bound to the DNA is also significantly populated. The high affinity for ssDNA is reflected in the total of ∼2100 A^2^ of buried surface across the entire interface. Consistent with the lack of symmetry within the trimer, the ssDNA binding interfaces are not identical for each protomer. The ability of the protomers to adapt to bind the substrate, and the fact that on average only ∼5 nucleotides are bound by each, provides a potential explanation for why a substantial population of RADX tetramers bound to dT25 is also observed in addition to the dominant trimers. The interaction with ssDNA is driven to a large extent by a large positively charged surface in OB2 that complements the ssDNA polyanion (Fig. 5b). Importantly, there are additional contacts with the ssDNA that are made by residues in one loop of the OB3 domain. Consistent with the asymmetry in the structure, the engagement with OB3 varies across the three protomers of the trimer with the OB3 loop from protomers A and B interacting with the ssDNA, but not the OB3 loop in protomer C. Correspondingly, the ssDNA binding interfaces of the protomers are not identical: protomers A and B each contact 6 nucleotides and bury ∼750 Å ^2^ of exposed surface whereas protomer C contacts 5 nucleotides and buries only ∼600 Å ^2^ due to the absence of the contribution from the OB3 loop (Table 1).

**Figure 5.**
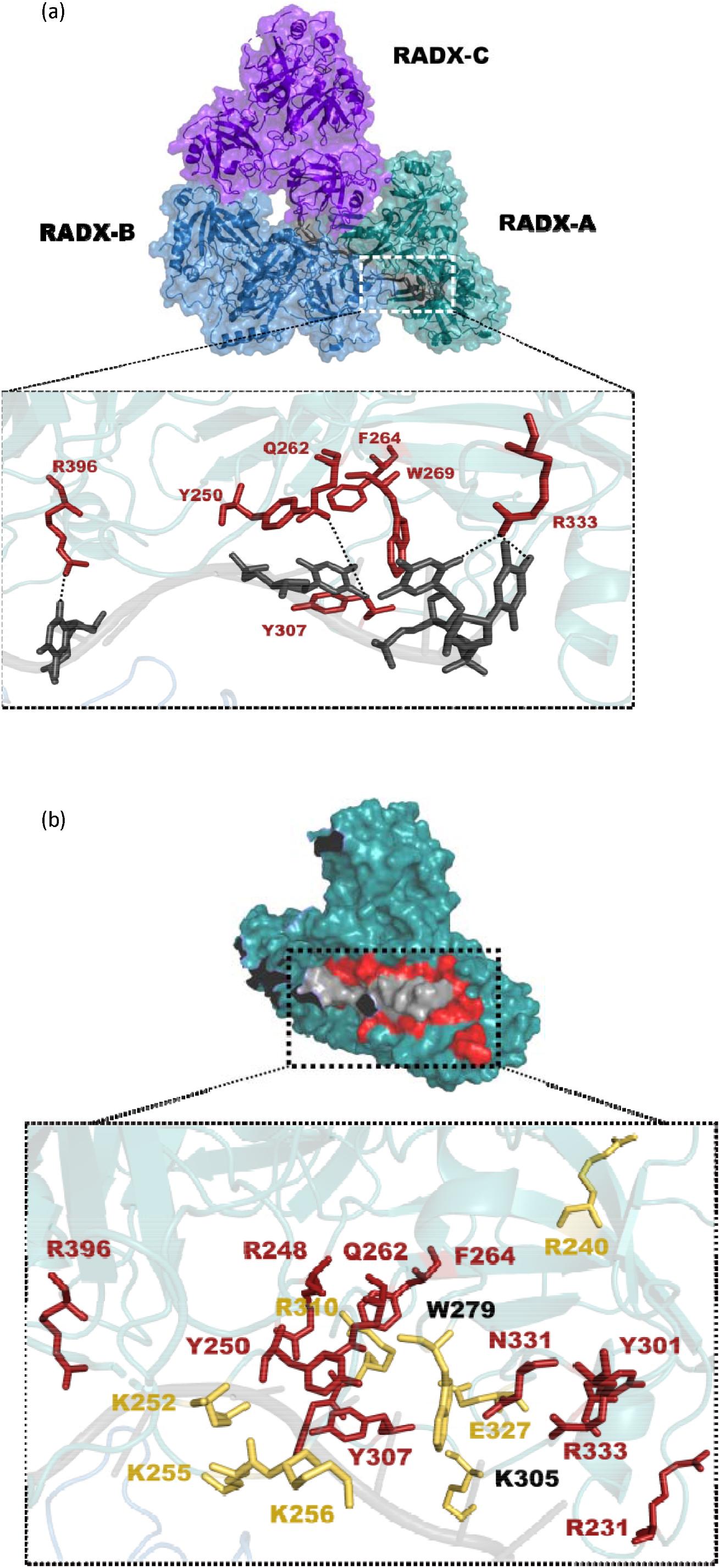
RADX binding of ssDNA is coupled to oligomerization. (a) The DNA binding site of RADX-A. The inset shows the residues involved in hydrogen bonding or pi-stacking interactions (in red) with the DNA (dark grey). H-bonds are shown as black dashed lines. (b) The full DNA interaction surface is shown in red in a surface representation of RADX-A (teal) with bound DNA (grey). The inset shows residues involved in hydrogen bonding or pi-stacking in any of the three RADX protomers (red). The yellow residues indicate the sites of mutation for the DNA binding Ob2M mutant, and the mutation site is seen to overlap only partially with the DNA binding site as shown by the two residues labelled in black (W279, K305).

**Table 1.** Analyses of the DNA binding surfaces of the three protomers in the RADX trimer.

| Protomer | Interface area (Å<sup>2</sup>) | *N<sub>SB</sub>+N<sub>HB</sub>+N<sub>pi</sub> | ΔG (kcal/mol) | Residues involved |
| --- | --- | --- | --- | --- |
| RADX-A | 744.5 | 10 | -11.5 | Y250, <u>Q262</u>, F264, W279, Y307, F309, <u>R333</u>, <u>R396</u> |
| RADX-B | 753.4 | 9 | -7.8 | <u>R248</u>, Y250, F264, W279, Y301, <u>K304</u>, Y307, F309 |
| RADX-C | 579.6 | 9 | -8.4 | <u>R232</u>, <u>R248</u>, Y250, <u>Q262</u>, F264, W279, Y307, <u>N331</u> |

\* Number of salt bridges + number of hydrogen bonds + number of pi-stacking interactions.
Underlined residues participate in hydrogen bonds.

The RADX-DNA interaction is mediated by both hydrogen bonds and pi-stacking interactions with aromatic residues in the DNA binding site. While the aromatic residues at the interaction interface such as W279, Y307 and F309 consistently participate in pi-stacking interactions, the identity and number of the hydrogen-bonding residues varies from protomer to protomer. For example, Q262, R333 and R396 in protomer A form hydrogen bonds with the ssDNA, whereas the residues involved in hydrogen bonding in protomer B are R248 and K304 (Table 1 and Fig. S5). To obtain additional insights, the free energy of RADX binding ssDNA was estimated using PDBePISA. The predictions ranged between −8 and −12 kcal/mol for the three different RADX protomers, much higher than the values of −1 to −2 kcal/mol determined for contributions to oligomerization. The higher values are fully consistent with the proposal that binding of DNA is coupled to RADX oligomerization and with the observation in the mass photometry data that binding to ssDNA drives formation of discrete oligomer populations (i.e. trimers and tetramers) (Fig. 1).

RADX variants to probe the DNA binding site(s) were designed previously based on sequence homology to RPA70, structural predictions and biochemical experiments. Two DNA-binding deficient variants containing multiple mutations in the predicted DNA binding surface of the OB2 domain were generated: one with nine single site mutations (R240E, R248E, K252E, K255E, K256E, W279A, K304E, R310E, E327A)^10,12^, and the other with only the K304A and E327A^13^. Both variants attenuated but did not abrogate DNA binding, which was interpreted as indicating other RADX domains must be involved in binding DNA. With the availability of the structure, we can now see that only two of the eight mutated residues (W279 and K304) directly contact the DNA (Fig. 5b). This explains the observation that ssDNA binding by these variants was reduced but not eliminated.

### RADX is unique but well conserved

RADX does not share significant sequence identity with proteins apart from other RADX orthologs. The closest match is to the large subunit of RPA (RPA70), although there is only 20% sequence identity between them. In contrast, the protein sequence is well conserved among RADX orthologs. Multiple sequence alignment shows that human RADX has a sequence identity of 75-100% with mammalian RADX orthologs and at least ∼50% sequence identity with RADX orthologs outside the *Mammalia* family (Fig. S6a S6b). The secondary structure elements present in the four domains of human RADX are seen to be well conserved across all the families where RADX is present. The highest variability is in the unstructured loop of human RADX (Fig. S7). The availability of an experimental structure provides a window to view sequence conservation in a structural and functional context and enables a search for structural homologs with similar folds. We were particularly interested in investigating if sequence conservation analysis would reveal residues in RADX whose structural and/or functional importance had yet to be investigated. Sequence conservation was assessed using the ConSurf^39^ server, which revealed as expected the structured regions of OB1, OB2 and OB3 are highly conserved (Fig. 6a). The key hydrogen-bonding residues involved in the oligomerization of RADX are also shown to be highly conserved, as are a majority of the residues of the DNA binding interface. There are two exceptions, F309 and R232, that are not conserved, the origin of which is not clear. In contrast to OB1, OB2 and OB3, OB4 has significant regions of variability, particularly in the large unstructured loop (S566-I676) inserted within the globular OB-fold domain (Fig. 6b). Multiple serine and threonine residues (e.g., T576, S584, T585 and S586) in this large loop are particularly intriguing as they are potential phosphorylation sites that could presumably modulate RADX activity.

**Figure 6.**
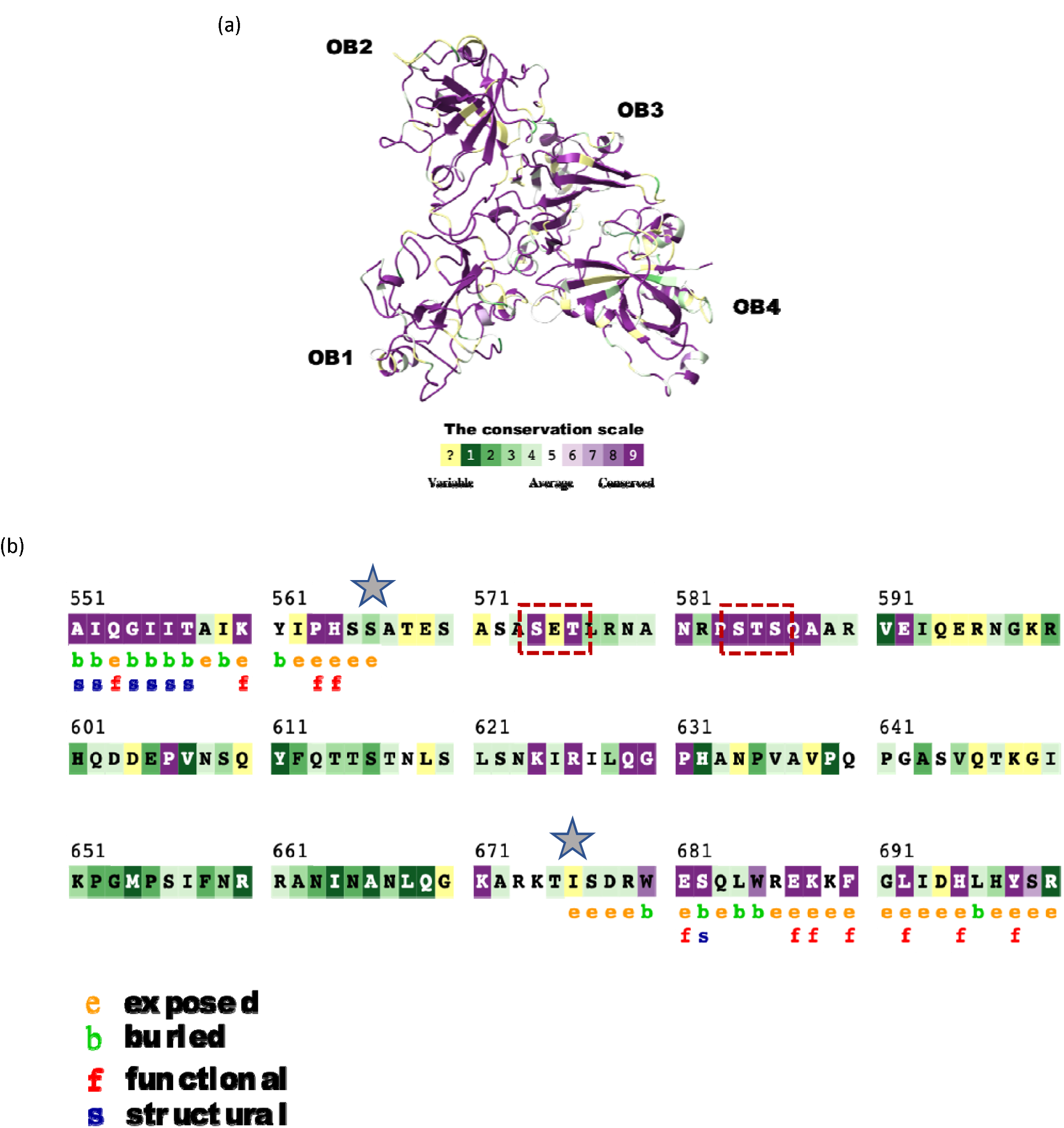
RADX is unique but well conserved. (a) Analysis of sequence conservation by ConSurf plotted onto the structure of RADX. The color code of the conservation scale is given below the structure. Regions of the protein with well-defined secondary structure are conserved. (b) Analysis of sequence conservation by ConSurf focused on the large loop in OB4 (S566-I676), delineated by grey stars. The loop has very poor sequence conservation, apart from small regions which include Ser and Thr residues, highlighted within red dashed boxes. (c) The high scored of 12.1 from the Dali server suggests RADX OB1 (blue)is a homolog of RPA70N domain (pink). Their structural similarity is reflected in the RMSD over all Ca atoms of 2.5 Å.

### RADX modulation of RAD51 filaments correlates with end-binding

We have shown previously that binding of ssDNA and interaction with RAD51 are required for RADX to inhibit RAD51 nucleoprotein filament formation on ssDNA *in vitro*^12^. We also demonstrated that RADX reduces the length of RAD51 filaments. Potential models for how RADX alters RAD51 filament formation include acceleration of the ATP hydrolysis rate of RAD51, sequestration of the ssDNA by RADX to prevent RAD51 binding or a combination of the two. To further explore the inhibition mechanism, define the RAD51 interaction interface, and address how RADX modulates RAD51 filaments, we turned again to electron microscopy.

In our previous study of RADX modifications of RAD51 filaments monitored by negative stain EM^12^, we were unable to locate RADX molecules due to their small size and the limited amount of RADX in the specimen. To overcome this limitation, an anti-RADX antibody attached to a 20 nm gold nanoparticle was used as a fiducial marker to locate RADX molecules, an approach used previously to locate BRCA2 bound to RAD51 filaments^45^. The negative stain EM micrographs obtained with nanogold labeling revealed RADX is binding to one end of RAD51 filaments (Fig. 7a left). Armed with this knowledge, negative stain EM data were collected on RADX bound to filaments without the gold nanoparticles. These data could then be analyzed by leveraging the availability of the high-resolution structure of RADX, allowing us to build low-resolution templates and find RADX more effectively in micrographs. The resulting particles from template-picked micrographs were assigned into 2D classes, which included free RAD51 filaments, free RADX molecules and RADX bound to the ends of RAD51 filaments (Fig. 7a right). The RADX bound filaments were distinguished on the basis of volumes and comparisons to previous structures of RAD51 filaments^46^. All 2D classes containing RAD51 filaments were isolated and grouped for ab-initio reconstruction and 3D classification with ∼4000 particles. This led to the generation of two low-resolution reconstructions at ∼20 Å, one containing only a RAD51 filament and the second in which the RAD51 filament is clearly visible with the additional unique volume at the end assigned to RADX (Fig. 7b). These results imply that RADX functions by binding to the ends of growing RAD51 filaments and capping further growth.

**Figure 7.**
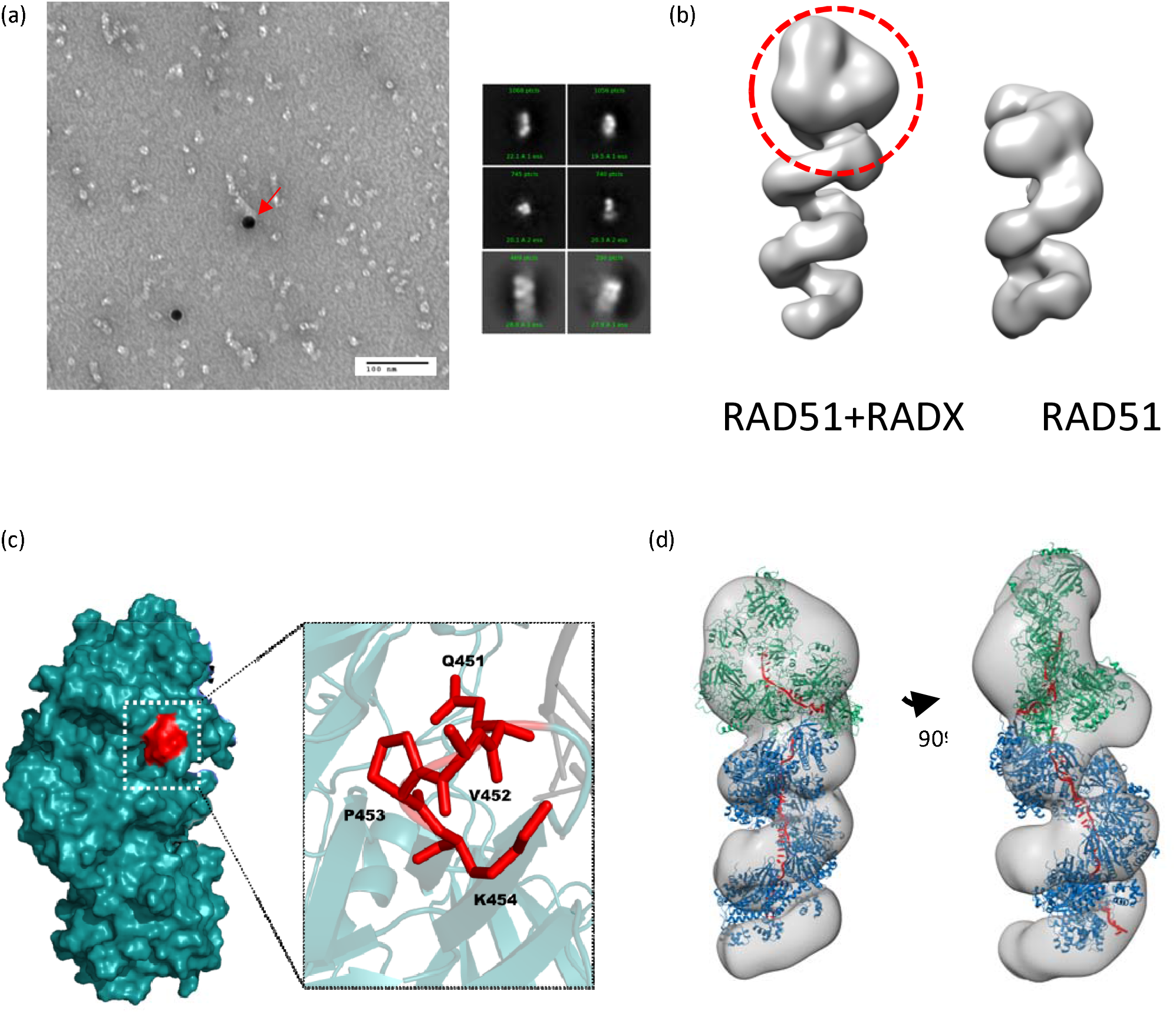
RADX modulation of RAD51 filaments correlates with end-binding. (a) (left) Negative stain EM micrograph showing anti-RADX Ab bound gold nanoparticles localized at the ends of RAD51 filaments. (right) 2D class averages of dataset containing RADX crosslinked to RAD51 filaments reveal populations of free RADX and RADX bound to RAD51 filaments. (b) 3D reconstructions of free RADX and RADX bound to RAD51 filaments. The extra volume assigned to RADX is circled in red. (c) The residues mutated in the RAD51 binding-deficient QVPK mutant displayed on the structure of RADX. These residues form a surface-exposed patch, consistent with the proposal that these are in RAD51 binding site. (d) Model of a RADX trimer bound to a RAD51 filament generated via manual docking. The RADX is shown in green, the RAD51 filament in blue (8BQ2) and the ssDNA in both models is shown in red.

RADX variants to probe RAD51 binding have been designed based on homology modeling and biochemical analysis. The RAD51 binding-deficient variant of RADX contained alanine substitutions for four consecutive residues in the OB3 domain (Q451-K454). In the high-resolution structure of the trimer, these residues are in a loop region and surface exposed (Fig. 7c), consistent with participation in a protein-protein interaction. This RADX QVPK variant led to elevated levels of the DNA-damage marker protein γH2AX in U2OS cells, and was unable to rescue slow rates of replication fork elongation in RADXΔ cells^12^. These observations show that the RAD51-RADX interaction is essential to maintain replication fork stability in cells.

To gain further insight into the RAD51-RADX interaction, we generated a model of a RAD51 filament with a RADX trimer bound to one end by docking the structures manually or with HADDOCK^34^ (Fig. 7d). The active residues specified for docking were the previously identified Q451-K454 for RADX, while CPORT was used to predict RAD51 active residues since no experimental information is available. Multiple HADDOCK-generated models and the manually docked model both fit remarkably well into our 3D reconstruction of the RAD51-RADX complex as reflected in the value of ∼0.9 for the goodness of fit assessed using the correlation coefficient from Chimera. The docked model shows the OB3 domain of RADX-B interacting with the RAD51 molecule on the 3’ end of the filament. The DNA binding site of RADX-A OB2 is remarkably well set up in a position to bind free ssDNA exposed at the end of the RAD51 filament. The model of RADX binding to RAD51 filaments on the 3’ end was chosen over the alternative model of 5’-end binding as we know that *in-vitro* RAD51 filaments on ssDNA extend preferentially in the 5’-3’ direction^47^, and so capping on the 3’ end would inhibit filament extension. More detailed analysis of the RAD51-RADX interface would be of great interest, but was not pursued from this model due to the low resolution of the negative stain EM reconstructions and the scarcity of information about the RADX interaction surface on RAD51.

## Discussion

RADX is required to maintain replication fork stability and is a direct regulator of RAD51 that counters the effect of positive regulators such as BRCA2. Knowledge of its structure and mechanism of action is of significance because tight regulation of RAD51 is needed to maintain genome stability. Thus, the high resolution cryo-EM structure of RADX opens up new horizons for exploring its function(s) as a modulator of RAD51 nucleofilament formation. RADX is structurally unique, with no matching protein folds in structural databases. Knowledge of the interfaces mediating oligomerization and DNA binding, energetically coupled and crucial to its function, facilitates precise design of separation of function mutants, which will enable more accurate definition of the roles of RADX in fork stalling, reversal and stabilization.

The absence of density for the large 111 residue loop (S566-I676) inserted within the OB4 domain strongly suggests it is disordered, consistent with predictions based on sequence. Hence, the functional role(s) of this loop remains a mystery. It is possible that the loop mediates interactions with as-yet-unknown binding partners. Alternatively, it may function in regulation of RADX activity. For example, there are number of highly conserved residues in the loop, including multiple serine and threonine residues that are potential sites of phosphorylation. Poly-phosphorylation has been shown to serve as a DNA mimic that can block DNA binding sites^48,49^, and could play such a role in RADX. Further biochemical and functional analysis of corresponding RADX loop mutants are required to test this hypothesis.

We have determined that RADX oligomerizes in a concentration and ssDNA-dependent manner, and that DNA binding drives the equilibrium to the trimeric and tetrameric states. Basal cell expression levels of RADX are in the nanomolar concentration range, where mass photometry shows RADX preferential formation of trimers, supporting the idea that this is a functionally relevant state. Mass photometry also shows that the preference for trimers remains independent of the length of ssDNA and that trimer particles form the majority population in cryo-EM micrographs. The highest masses observed for RADX on dT25, dT40 and dT60 correspond to a tetramer, pentamer and septamer, respectively. Thus, RADX oligomerization does not scale linearly with the increase in substrate length. Since mass photometry only reports on the total mass of a complex, if RADX formed single filaments, the oligomer would grow by one protomer for each 5-6 nucleotides in additional length and one would expect a predominance of septamers and undecamers for dT40 and dT60, respectively, but this is not observed. Rather, the results for dT40 and dT60 are consistent only with combinations of discrete lower order dimers, trimers and tetramers spaced apart on the substrate, not a single large oligomer. The results also align with the DNA footprint for polydT of 19-27 residues, a range that corresponds to trimers and tetramers and not higher order RADX-DNA oligomers.

Figure 8 shows a diagrammatic representation of our model of how RADX functions. Upon exposure of ssDNA at stalled replication forks, RPA binds and protects the DNA from degradation, and recruits a range of damage repair factors. RPA is replaced by RAD51 via the action of a mediator such as BRCA2. RADX is localized to the vicinity of RAD51 through its protein interaction surface and can bind exposed ssDNA at the termini of the expanding RAD51 filament. RADX binding to ssDNA is energetically coupled to oligomerization and forms stable caps at the end of the filament, effectively blocking further extension. We have previously shown that the rate of ATP hydrolysis by RAD51 increases in the presence of RADX, and that binding of BRCA2 to RAD51 slows the ATP hydrolysis rate. Therefore, the mechanism by which RADX disassembles RAD51 filaments is likely to consist of two parts: end capping of the RAD51 filament by RADX oligomers to block extension, and a conformational change in RAD51 upon interacting with the terminal RADX that accelerates ATP hydrolysis and therefore release of RAD51 from DNA.

**Figure 8.**
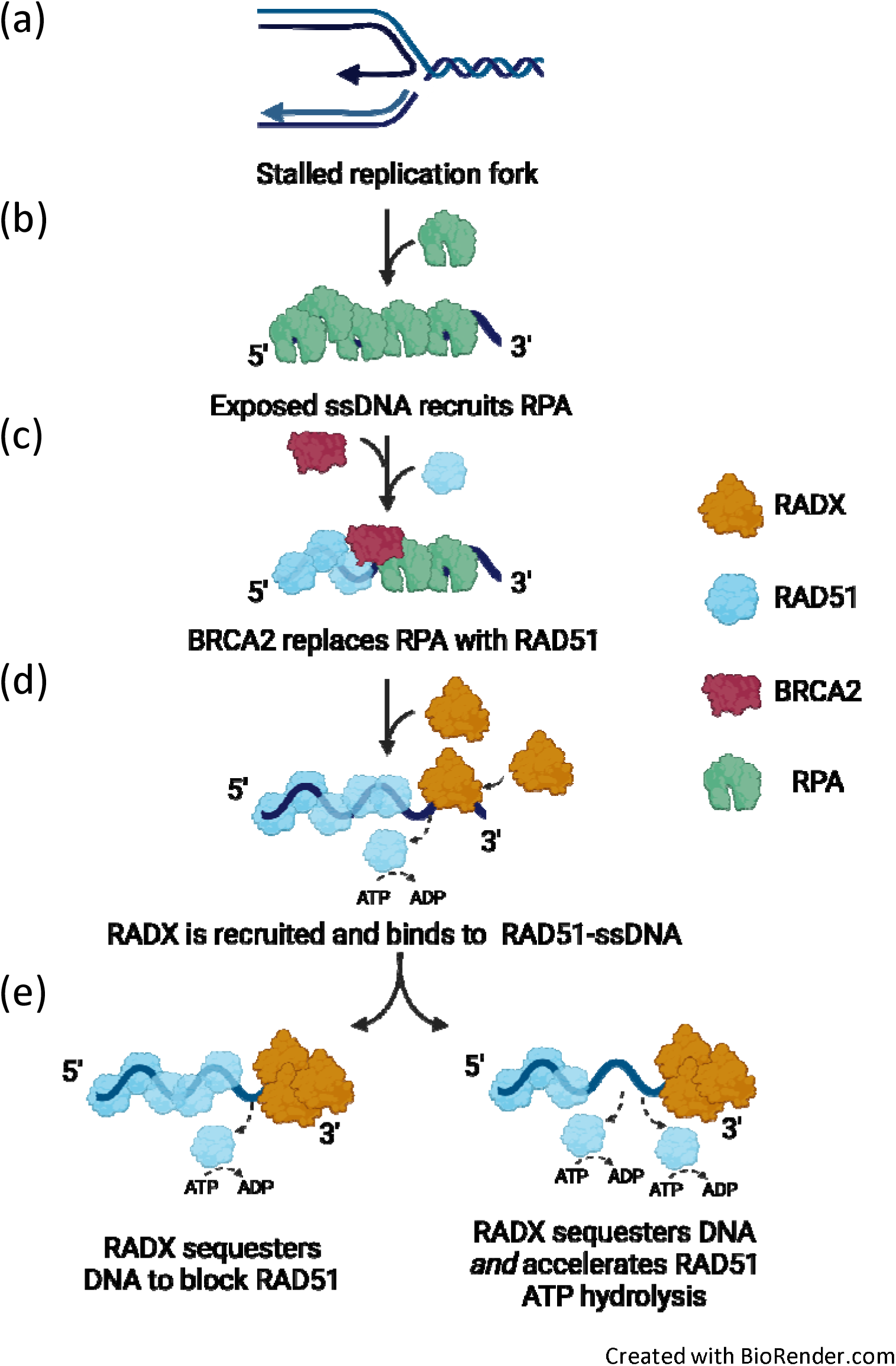
Model for the mechanism of RADX action. Upon replication fork stalling, single stranded DNA is exposed (a), which is then bound by RPA to protect the ssDNA from damaging agents (b). RPA is replaced on ssDNA with RAD51 by the action of BRCA2, a mediator protein (c). RAD51 binds cooperatively to form filaments on ssDNA. RADX is recruited to ssDNA where it binds the ends of the growing RAD51 filament (d). It promotes filament disassembly by either i) blocking expansion at the end of the filament as RAD51 hydrolyses ATP and detaches from ssDNA or ii) both blocking filament expansion and accelerating the ATP hydrolysis rate of RAD51(e).

The model places RADX at the 3’ end of the filament based on RADX interacting with RAD51 filaments in isolation. At a stalled replication fork in the cell, other modulators such as BRCA2 and the RAD51 paralog proteins will also be present^5,50^. While BRCA2 is seen as primarily a RAD51 filament nucleation factor, the RAD51 paralog proteins bind the 5’ end of the filament and stabilize it, promoting 3’-5’ filament extension^9^. It is therefore possible that RADX caps either or both the 3’ and the 5’ ends of the RAD51 nucleoprotein filament depending on the other proteins acting at the replication fork at that time.

Filament formation by RAD51 is a co-operative process^51^, with the rate of growth of filaments on ssDNA measured as 3 orders of magnitude higher than the nucleation rate. Regulators like BRCA2 play a crucial role during nucleation, first by making DNA available by removing RPA and then stabilizing an oligomer of RAD51 on ssDNA. Once filament formation has begun, the high cooperativity results in rapid elongation of the filament, which would limit the efficacy of an inhibition mechanism that relies solely on out-competing RAD51 for ssDNA. A mechanism that physically blocks RAD51 and shifts the equilibrium of DNA binding to disassembly is both plausible and supported by *in-vitro* experimental evidence, including the disassembly of RAD51 filament by RADX and corresponding increase in the hydrolysis rate of RAD51 bound ATP^12^, as well as the localization of RADX at the termini of filaments shown in this study. Additional functional studies using separation of function mutants designed using the now-available RADX structure as well as determination of a high-resolution structure of RADX bound to RAD51 will test and refine this proposed mechanism of RADX function.

## Materials and methods

### Expression and purification of RADX

RADX and a C-terminal truncation construct (RADX 43-560, RADX-N) were expressed and purified from three different expression systems, *E.coli*, High-five insect cells (Expression Systems) and Expi293F mammalian cells (Thermo Fisher). RADX and RADX-N were expressed in *E.coli* BL21(DE3) Star cells using a pIF332 plasmid construct with MBP-tagged RADX^14^. Cells were transformed with plasmid and then allowed to grow to 37 °C until the OD600 was 1. They were then induced with 1 mM IPTG and grown for 12-16 h at 18 °C, then cells were harvested by centrifugation. RADX was expressed in Sf9 insect cells using baculovirus generated from a 438-C plasmid construct with RADX fused to a HRV-3C protease-cleavable N-terminal 6xHis-MBP tag. Cells were infected with baculovirus at MOI of 1 at 140 rpm for 70 h at 27 °C. Similarly, RADX-N was expressed in High-five insect cells infected with baculovirus at MOI of 1 for 45 h at 27 °C. Cells were harvested by centrifugation at 500xg for 10 min. RADX was expressed in human Expi293F cells using BacMam^16^ baculovirus generated from a pEZT-BM^17^ plasmid construct with RADX fused with a HRV-3C protease-cleavable N-terminal 6xHis-MBP and a C-terminal Flag tag. Cells were infected with BacMam baculovirus at MOI of 5 in the presence of 3 mM butyrate at 125 rpm, 37 °C, 8% CO2 for 48 h and harvested by centrifugation at 500xg for 10 min. Cell pellets were homogenized in 5 ml/g lysis buffer (20 mM Tris-HCl at pH 7.5, 150 mM NaCl, 1% Triton X-100, 10% glycerol, 10 mM NaH_2_PO_4_, 10 mM NaP_2_O_7_, 100 μM ZnCl_2_, 10 mM NaF, 1 Roche protease inhibitor tablet), lysed by sonication and the supernatant isolated by centrifugation. The protein was then affinity purified over amylose resin in a buffer containing 50 mM Tris-HCl at pH 7.5, 500 mM NaCl, 5% glycerol, 0.05% Tween-20, 0.3 mM PMSF (pheynlmethylsulphonylfluoride) and 2.5 mM BME (β-mercaptoethanol). The protein was eluted using 20 mM maltose and then further purified by size exclusion chromatography (SEC) using a 24 mL Superose 6 column (Cytiva). The purification protocol was very similar for protein expressed from High-five cells as well as mammalian cells. Depending on the downstream experiment, the following modifications were performed: for cryo-EM, an excess of dT25 ssDNA was added to the protein prior to SEC. For experiments requiring the protein to be DNA-free, DNase was added to the lysis buffer and the SEC step was replaced by another affinity chromatography using heparin resin. For mass photometry experiments the protein was purified without Tween-20.

### Expression and purification of RAD51

RAD51 was expressed in BLR(DE3)pLysS *E.coli* cells, a RecA deficient cell line. Competent cells were transformed with a pMBP-His-Tev-RAD51 plasmid and grown at 37 °C until the OD600 reached 0.6-0.8. The cells were then induced with 500 mM IPTG and grown overnight at 18 °C for expression. Cells were harvested by centrifugation, lysed by sonication after homogenization in lysis buffer (50 mM Tris-HCl at pH 7.5, 5 mM EDTA, 200 mM KCl, 1 mM DTT, 10% Sucrose, 0.01% NP-40 and 1 mM PMSF). The protein was purified by affinity chromatography using first amylose resin and then heparin resin. Protein was bound to amylose resin equilibrated with 20 mM Tris-HCl at pH 7.5, 0.5 mM ETDA, 100 mM KCl, 2.5 mM BME, 10% glycerol and 0.01% NP40, and eluted using 5 column volumes (CV) of 20 mM Maltose in the same buffer. The protein was then cleaved using TEV protease, loaded onto a heparin column equilibrated with the buffer above and eluted with a gradient over 12 CV to 1 M KCl.

### Mass photometry

Mass photometry studies were performed using a TwoMP system from Refeyn^18^. All samples were run in 1X PBS (Gibco, Thermo 10010023). The mass calibrations were performed using standard curves generated using gamma-globulin, thyroglobulin and beta-amylase (20-50 nM concentrations for each). RADX was diluted using 50 mM Tris-HCl at pH 7.5, 100 mM NaCl, 2% Glycerol, 2.5 mM BME to a concentration of 425 nM for the 50 nM readings and 850 nM for the 100 nM readings. The samples with DNA had 1:1 ratios of the specified DNA oligomer added 15-30 min before data collection. All samples were run by setting focus with 15 μL of PBS and then adding 2 μL of the sample for data collection. Data was collected using AcquireMP 2.3 and analyzed using DiscoverMP 2.3. Plots were exported to and formatted in Inkscape.

### Steady state rotational anisotropy and fluorescence quenching measurements

Steady state fluorescence depolarization (rotational anisotropy) was used to measure the affinity of RADX for various lengths of fluorescein-labelled ssDNA. Samples were prepared as 25 μl solutions containing 5 nM fluorescein-labelled ssDNA in binding buffer: 20 mM Tris at pH 7.5, 150 mM NaCl, 1 mM DTT, 5% glycerol. RADX was titrated over the range 0-160 nM until saturation was reached. A filter-based Synergy Neo2 plate reader (Biotek) was used to collect data and calculate anisotropy. Samples were excited with vertically polarized light at 485/20 nm and vertical and horizontal emissions were measured at 530/25 nm with a 510 nm dichroic mirror. Apparent dissociation constants (*K*_d_) were obtained by fitting to a sigmoidal curve using Prism.

RADX binding of a polydT substrate (Midland Certified Reagent Co) was monitored in binding buffer (20 mM Tris at pH 7.5, 100 or 200 mM NaCl, 1 mM DTT, 5% glycerol) under stoichiometric conditions. A constant concentration of RADX (100 nM) was titrated with increasing amounts of poly-dT. Binding was tracked with a Horiba Jobin Yvon Fluoromax-3 fluorometer. Data were collected at an excitation wavelength at 290 nm and emission wavelength at 340 nm at 25 °C. The occluded site size was determined by the intersection of the linear parts of the titration curve, plotted using Prism software. Measurements were corrected for dilution, photo-bleaching, and inner filter effects.

### SEC-SAXS data collection and analysis for RADX-dT25

Samples of RADX from mammalian as well as High-five cells were prepared for SEC-SAXS. The protein was purified as detailed above, bound to dT25 and then concentrated to 20 −30 μM. The SAXS profiles for the RADX-dT25 complex were collected in SEC-SAXS mode at the ALS beamline 12.3.1 at the Lawrence Berkely National Laboratory in Berkeley, California^19^. The X-ray wavelength *λ* was 1.03 Å and the sample-to-detector distance was set to 1.5 m resulting in scattering vectors, *q*, ranging from 0.01 to 0.5 Å^−1^. All experiments were performed at 20 °C and data were processed as described elsewhere^20,21^. Briefly, the flow-through SAXS cell was directly coupled with an online Agilent 1260 Infinity HPLC system using a Shodex KW803 column (Shodex™). The column was equilibrated with running buffer (20 mM Tris at pH 7.5, 150 mM NaCl, 2% glycerol, 0.05% Tween-20) with a flow rate of 0.5 ml/min. A 50 μL sample was run through the SEC and 3.0 s X-ray exposures were collected continuously during a ∼35 min elution. The SAXS frames recorded prior to the protein elution peak were used as buffer blanks to subtract from all other frames. The programs SCÅTTER 3.1 and RAW^22,23^ were used to analyse the data and determine the *Rg*, *Dmax* and *P(r)* distribution. Protein purified from High-five cells was seen to produce better quality SEC-SAXS data, and only the results for these data are described and discussed below.

### Cryo-EM sample preparation and data collection

Samples for cryo-EM were prepared as follows: RADX was incubated with a molar excess of dT25 and then purified by SEC using a 24 mL Superose 6 column. The final grids used to generate the high-resolution structures were prepared by taking a 20 μL aliquot adjusted to 1 μM RADX then incubating with 10 μM BS3 for 2 hours on ice. Glow-discharged holey-carbon R 1.2/1.3 300 mesh copper grids were prepared, and 2.5 μL of sample was applied and blotted away using filter paper (Ted Pella) twice. The grid was immediately loaded onto a Vitrobot Mark IV at 4 °C and 100% humidity, sample was added again, grid was blotted and plunge-frozen in liquid ethane. Grids were screened using a Glacios 200 kV TEM (ThermoFisher) equipped with a Falcon4 direct electron detector. Large-scale datasets were collected using a Titan Krios G4 300 kV TEM (ThermoFisher) equipped with a Gatan K3 BioQuantum direct electron detector functioning in counting mode. Each dataset was collected in a single session from a single grid at a magnification of 105 kx (0.82 Å /pixel) and a dose rate of 16.061 e/pix/second over vacuum. The total exposure time per micrograph was 2.3 seconds and contained 50 frames, making the exposure time per frame 50 milliseconds. A total of 15072 images were collected using EPU (ThermoFisher) in fast acquisition mode with a defocus range of −2.0 to −0.8 μm.

### Cryo-EM data processing and analysis

Data processing and analysis was performed using cryoSPARC^24^. Movies were imported, patch motion correction and patch CTF estimation were run as pre-processing. The micrographs were first blob-picked, particles were extracted with down-sampling from 512 to 128 pixels (px). After several rounds of 2D classification to eliminate junk particles, a varied selection of classes were chosen as templates. A small subset of 200 micrographs were template picked and the extracted 21,129 particles were input into Topaz^25^ to train a model. This model was then used to pick the entire dataset. The resulting particles were extracted at 512 px and subjected to several rounds of 2D classification, during which trimer and tetramer particles were separated. The trimer and tetramer particle sets contained 800,000 and 300,000 particles, respectively. Each particle set was then put through *ab-initio* reconstruction and several rounds of 3D classification followed by non-uniform refinement for each class. The particle sets with highest resolution after refinement were used for further processing. In the case of the trimer, the final particle set of 158,000 particles was cleaned with particle subtraction and then local refinement was used focusing on each dimer pair in the trimer to generate a composite map of resolution 3 Å. For the tetramer, local refinement focusing on the fourth RADX was used to generate the final map.

An initial model based on the map was generated by ModelAngelo^26^, as the lack homology model meant a *de novo* structure needed to be generated from the map. This initial model was then further refined using both the real space refine package and secondary structure/stereochemical restraints in Phenix^27^ and visualized using Coot^28^. The model was validated using MolProbity^29^. Figures were generated using Chimera^30^, ChimeraX^31^ and Pymol^32^. All programs were run within the SBGrid platform^33^.

### RADX localization and negative-stain EM studies

To determine localization of RADX on RAD51 nucleoprotein filaments, 20 nm gold nanoparticles from an antibody conjugation kit (Abcam Ab188215) were conjugated using the kit protocol to an anti-RADX mouse monoclonal Ab (CXorf57, Santa Cruz Biotechnology, sc-514563). The conjugated antibody was then incubated with an excess of RADX for 30 min on ice. RAD51 filaments were formed by adding RAD51 (8 μM), ATP (5 mM) and dT72 (6 μM) in a buffer containing 25 mM HEPES at pH 7.5, 25 mM KCl and 4 mM MgCl_2_. The filament mix was incubated at 37 °C for 15 min, then RADX-Ab-AuNP was added to a final concentration of 80 nM (1:100 RADX:RAD51). The mix was immediately fixed on carbon coated 400 mesh copper grids (CF-400-Cu, EMS), stained with uranyl formate and imaged using a Morgagni 100 kV electron microscope equipped with a 1kX1k CCD camera.

To collect negative stain data on RADX bound to RAD51 nucleofilaments in the absence of gold nanoparticles, the same protocol as above was followed to generate the RAD51 filaments, and RADX was also added as above. BS3 cross-linker was also added to the mix to a final concentration of 1 mM immediately after the addition of RADX, and the reaction was then kept at room temperature for 30 min. The reaction was quenched with Tris-HCl at pH 8.0 to a final concentration of 50 mM, and the sample was fixed to a CF-400-Cu as above. Data were collected on an FEI TF20 200 kV TEM with a 4kX4K CCD camera. These data were processed using cryoSPARC.

### Computational analyses to determine sequence conservation, structural homology and docking of RADX and RAD51

The PDBePISA server was used to analyze the residues involved at the interaction interfaces between RADX protomers, as well as interaction interfaces between RADX and ssDNA. The estimates of the energetics of these interactions were obtained from the same server. For HADDOCK^34^ docking, the structure of RAD51 in filament form was obtained from the PDB (8BQ2). Both RADX trimer and tetramer structures were used as docking partners, with the residues Q451-Y457 used as active residues on RADX. As the binding site of RADX on RAD51 is unknown, the interaction site was predicted using CPORT^35^. The active residues in RAD51 used for docking were: Y54, A79, K80, P83-S97, I99, I100, E118, G120, F129, R130, D257, E258, L303, R304 and P318. Docking was performed using default parameters and the top 10 models were compared to the molecular envelope to find the best fit. For manual docking the RADX trimer and the RAD51 filament were docked using the ‘Fit to map’ function of Chimera, and then realigned based on the known interaction surfaces.

Sequence-based homology and alignment were analyzed using BLAST^36^ and ALIGN^37^ on the UniProt^38^ website. The UniProt sequence Q6SNI4-1 was used as the query sequence in each case. A BLAST search of the top 250 sequences showed 166 of the 250 sequences belonged to the taxonomic class Mammalia, and 84 belonging to the Euteleostomi clade but not to Mammalia. ALIGN was run on sequence subsets of all mammalian sequences aligned to the query sequence and all non-mammalian sequences aligned to the query sequence. Sequence conservation was assessed using ConSurf^39^. Due to the absence of the large S566-I676 loop in the experimental structure, data were generated with just the structure as input and with a combination of the structure and multiple sequence alignment (MSA) inputs to cover the entire protein. Both sets of data showed identical conservation levels for the residues present in the structure. Analysis of similarity to known structures was performed on the DALI^40^ and FoldSeek^41^ servers using the PDB25 search for DALI and PDB100 search for FoldSeek. Both programs gave very similar results, and results from DALI were used for further analysis.

## Acknowledgements

We thank Prof. Melanie Ohi, Dr. Jason Porta, Dr. Heather Kroh, Dr. Elad Binshtin and Dr. Elwood Mullins for advice and assistance with cryo-EM data analysis. Dr. Carl Schiltz provided advice on optimization of protein cross-linking. This work was supported by grants from the US National Institutes of Health: R35 GM118089 (WJC); P01 CA092584 (MST, DC, WJC), R01 GM116616 (DC). The EM data were acquired at the Vanderbilt Center for Structural Biology Cryo-EM Facility with the support of NIH grant S10 OD030292 for the purchase of the Glacios cryo-TEM.

## Data availability

Structural data has been deposited to the PDB and EMDB. This includes structural co-ordinates, cryo-EM maps, and details of data collection. Accession codes for the RADX trimer are PDB ID 8U5Y/EMD ID 41939 and tetramer PDB ID 8U61/EMD ID 41940. All other data are available in this paper.

## References

(1) Berti, M., Cortez, D., and Lopes, M. (2020) The plasticity of DNA replication forks in response to clinically relevant genotoxic stress. Nat. Rev. Mol. Cell Biol. 21, 633–651.

(2) Cortez, D. (2019) Replication-Coupled DNA Repair. Mol. Cell 74, 866–876.

(3) Zellweger, R., Dalcher, D., Mutreja, K., Berti, M., Schmid, J. A., Herrador, R., Vindigni, A., and Lopes, M. (2015) Rad51-mediated replication fork reversal is a global response to genotoxic treatments in human cells. J. Cell Biol. 208, 563–579.

(4) Hashimoto, Y., Chaudhuri, A. R., Lopes, M., and Costanzo, V. (2010) Rad51 protects nascent DNA from Mre11-dependent degradation and promotes continuous DNA synthesis. Nat. Struct. Mol. Biol. 17, 1305–1311.

(5) Schlacher, K., Christ, N., Siaud, N., Egashira, A., Wu, H., and Jasin, M. (2011) Double-strand break repair-independent role for BRCA2 in blocking stalled replication fork degradation by MRE11. Cell 145, 529–542.

(6) Sarbajna, S., and West, S. C. (2014) Holliday junction processing enzymes as guardians of genome stability. Trends Biochem. Sci. 39, 409–419.

(7) Kolinjivadi, A. M., Sannino, V., de Antoni, A., Técher, H., Baldi, G., and Costanzo, V. (2017) Moonlighting at replication forks – a new life for homologous recombination proteins BRCA1, BRCA2 and RAD51. FEBS Lett. 591, 1083–1100.

(8) Sullivan, M. R., and Bernstein, K. A. (2018) RAD-ical new insights into RAD51 regulation. Genes (Basel). 9.

(9) Taylor, M. R. G., Špírek, M., Jian Ma, C., Carzaniga, R., Takaki, T., Collinson, L. M., Greene, E. C., Krejci, L., and Boulton, S. J. (2016) A Polar and Nucleotide-Dependent Mechanism of Action for RAD51 Paralogs in RAD51 Filament Remodeling. Mol. Cell 64, 926–939.

(10) Dungrawala, H., Bhat, K. P., Le Meur, R., Chazin, W. J., Ding, X., Sharan, S. K., Wessel, S. R., Sathe, A. A., Zhao, R., and Cortez, D. (2017) RADX Promotes Genome Stability and Modulates Chemosensitivity by Regulating RAD51 at Replication Forks. Mol. Cell 67, 374–386.e5.

(11) Bhat, K. P., Krishnamoorthy, A., Dungrawala, H., Garcin, E. B., Modesti, M., and Cortez, D. (2018) RADX Modulates RAD51 Activity to Control Replication Fork Protection. Cell Rep. 24, 538–545.

(12) Adolph, M. B., Mohamed, T. M., Balakrishnan, S., Xue, C., Morati, F., Modesti, M., Greene, E. C., Chazin, W. J., and Cortez, D. (2021) RADX controls RAD51 filament dynamics to regulate replication fork stability. Mol. Cell 81, 1074–1083.e5.

(13) Schubert, L., Ho, T., Hoffmann, S., Haahr, P., Guérillon, C., and Mailand, N. (2017) RADX interacts with single-stranded DNA to promote replication fork stability. EMBO Rep. 18, 1991–2003.

(14) Zhang, H., Schaub, J. M., and Finkelstein, I. J. (2020) RADX condenses single-stranded DNA to antagonize RAD51 loading. Nucleic Acids Res. 48, 7834–7843.

(15) Mohamed, T., Adolph, M. B., and Cortez, D. (2022) Oligomerization of DNA replication regulatory protein RADX is essential to maintain replication fork stability. J. Biol. Chem. 298, 101672.

(16) Gradia, S. D., Ishida, J. P., Tsai, M.-S., Jeans, C., Tainer, J. A., and Fuss, J. O. (2017) MacroBac: New Technologies for Robust and Efficient Large-Scale Production of Recombinant Multiprotein Complexes. Methods Enzymol. 592, 1–26.

(17) Morales-Perez, C. L., Noviello, C. M., and Hibbs, R. E. (2016) Manipulation of Subunit Stoichiometry in Heteromeric Membrane Proteins. Structure 24, 797–805.

(18) Young, G., Hundt, N., Cole, D., Fineberg, A., Andrecka, J., Tyler, A., Olerinyova, A., Ansari, A., Marklund, E. G., Collier, M. P., Chandler, S. A., Tkachenko, O., Allen, J., Crispin, M., Billington, N., Takagi, Y., Sellers, J. R., Eichmann, C., Selenko, P., Frey, L., Riek, R., Galpin, M. R., Struwe, W. B., Benesch, J. L. P., and Kukura, P. (2018) Quantitative mass imaging of single molecules HHS Public Access. Science (80-.). 360, 423–427.

(19) Classen, S., Hura, G. L., Holton, J. M., Rambo, R. P., Rodic, I., McGuire, P. J., Dyer, K., Hammel, M., Meigs, G., Frankel, K. A., and Tainer, J. A. (2013) Implementation and performance of SIBYLS: a dual endstation small-angle X-ray scattering and macromolecular crystallography beamline at the Advanced Light Source. J. Appl. Crystallogr. 46, 1–13.

(20) Dyer, K. N., Hammel, M., Rambo, R. P., Tsutakawa, S. E., Rodic, I., Classen, S., Tainer, J. A., and Hura, G. L. (2014) High-Throughput SAXS for the Characterization of Biomolecules in Solution: A Practical Approach, in Structural Genomics: General Applications (Chen, Y. W., Ed.), pp 245–258. Humana Press, Totowa, NJ.

(21) Hura, G. L., Menon, A. L., Hammel, M., Rambo, R. P., Poole, F. L., Tsutakawa, S. E., Jenney, F. E., Classen, S., Frankel, K. A., Hopkins, R. C., Yang, S. J., Scott, J. W., Dillard, B. D., Adams, M. W. W., and Tainer, J. A. (2009) Robust, high-throughput solution structural analyses by small angle X-ray scattering (SAXS). Nat. Methods 6, 606–612.

(22) Nielsen, S. S., Toft, K. N., Snakenborg, D., Jeppesen, M. G., Jacobsen, J. K., Vestergaard, B., Kutter, J. P., and Arleth, L. (2009) BioXTAS RAW, a software program for high-throughput automated small-angle X-ray scattering data reduction and preliminary analysis. J. Appl. Crystallogr. 42, 959–964.

(23) Hopkins, J. B., Gillilan, R. E., and Skou, S. (2017) BioXTAS RAW: improvements to a free open-source program for small-angle X-ray scattering data reduction and analysis. J. Appl. Crystallogr. 50, 1545–1553.

(24) Punjani, A., Rubinstein, J. L., Fleet, D. J., and Brubaker, M. A. (2017) CryoSPARC: Algorithms for rapid unsupervised cryo-EM structure determination. Nat. Methods 14, 290–296.

(25) Bepler, T., Morin, A., Rapp, M., Brasch, J., Shapiro, L., Noble, A. J., and Berger, B. (2019) Positive-unlabeled convolutional neural networks for particle picking in cryo-electron micrographs. Nat. Methods 16, 1153–1160.

(26) Jamali, K., Kimanius, D., and Scheres, S. (2022) ModelAngelo: Automated Model Building in Cryo-EM Maps. arXiv:2210.00006 [q-bio.QM].

(27) Liebschner, D., Afonine, P. V, Baker, M. L., Bunkóczi, G., Chen, V. B., Croll, T. I., Hintze, B., Hung, L. W., Jain, S., McCoy, A. J., Moriarty, N. W., Oeffner, R. D., Poon, B. K., Prisant, M. G., Read, R. J., Richardson, J. S., Richardson, D. C., Sammito, M. D., Sobolev, O. V, Stockwell, D. H., Terwilliger, T. C., Urzhumtsev, A. G., Videau, L. L., Williams, C. J., and Adams, P. D. (2019) Macromolecular structure determination using X-rays, neutrons and electrons: recent developments in Phenix. Acta Crystallogr. Sect. D, Struct. Biol. 75, 861–877.

(28) Emsley, P., Lohkamp, B., Scott, W. G., and Cowtan, K. (2010) Features and development of Coot. Acta Crystallogr. Sect. D Biol. Crystallogr. 66, 486–501.

(29) Chen, V. B., Arendall, W. B. 3rd, Headd, J. J., Keedy, D. A., Immormino, R. M., Kapral, G. J., Murray, L. W., Richardson, J. S., and Richardson, D. C. (2010) MolProbity: all-atom structure validation for macromolecular crystallography. Acta Crystallogr. D. Biol. Crystallogr. 66, 12–21.

(30) Pettersen, E. F., Goddard, T. D., Huang, C. C., Couch, G. S., Greenblatt, D. M., Meng, E. C., and Ferrin, T. E. (2004) UCSF Chimera--a visualization system for exploratory research and analysis. J. Comput. Chem. 25, 1605–1612.

(31) Pettersen, E. F., Goddard, T. D., Huang, C. C., Meng, E. C., Couch, G. S., Croll, T. I., Morris, J. H., and Ferrin, T. E. (2021) UCSF ChimeraX: Structure visualization for researchers, educators, and developers. Protein Sci. 30, 70–82.

(32) Schrödinger, L. (2015) The PyMOL Molecular Graphics System, Version∼1.8.

(33) Morin, A., Eisenbraun, B., Key, J., Sanschagrin, P. C., Timony, M. A., Ottaviano, M., and Sliz, P. (2013) Cutting edge: Collaboration gets the most out of software. Elife 2.

(34) Dominguez, C., Boelens, R., and Bonvin, A. M. J. J. (2003) HADDOCK: A Protein−Protein Docking Approach Based on Biochemical or Biophysical Information. J. Am. Chem. Soc. 125, 1731–1737.

(35) de Vries, S. J., and Bonvin, A. M. J. J. (2011) CPORT: A Consensus Interface Predictor and Its Performance in Prediction-Driven Docking with HADDOCK. PLoS One 6, 1–12.

(36) Altschul, S. F., Gish, W., Miller, W., Myers, E. W., and Lipman, D. J. (1990) Basic local alignment search tool. J. Mol. Biol. 215, 403–410.

(37) Sievers, F., Wilm, A., Dineen, D., Gibson, T. J., Karplus, K., Li, W., Lopez, R., McWilliam, H., Remmert, M., Söding, J., Thompson, J. D., and Higgins, D. G. (2011) Fast, scalable generation of high-quality protein multiple sequence alignments using Clustal Omega. Mol. Syst. Biol. 7.

(38) Consortium, T. U. (2023) UniProt: the Universal Protein Knowledgebase in 2023 - Google Scholar 51, 523–531.

(39) Yariv, B., Yariv, E., Kessel, A., Masrati, G., Chorin, A. Ben, Martz, E., Mayrose, I., Pupko, T., and Ben-Tal, N. (2023) Using evolutionary data to make sense of macromolecules with a “face-lifted” ConSurf. Protein Sci. 32, 1–12.

(40) Holm, L. (2020) DALI and the persistence of protein shape. Protein Sci. 29, 128–140.

(41) Kempen, M. van, Kim, S. S., Tumescheit, C., Mirdita, M., Gilchrist, C. L. M., Söding, J., and Steinegger, M. (2022) Foldseek: fast and accurate protein structure search. bioRxiv 2022.02.07.479398.

(42) Theobald, D. L., Mitton-Fry, R. M., and Wuttke, D. S. (2003) Nucleic acid recognition by OB-fold proteins. Annu. Rev. Biophys. Biomol. Struct. 32, 115–133.

(43) Krissinel, E., and Henrick, K. (2007) Inference of Macromolecular Assemblies from Crystalline State. J. Mol. Biol. 372, 774–797.

(44) Brosey, C. A., Soss, S. E., Brooks, S., Yan, C., Ivanov, I., Dorai, K., and Chazin, W. J. (2015) Functional Dynamics in Replication Protein A DNA Binding and Protein Recruitment Domains. Structure 23, 1028–1038.

(45) Shahid, T., Soroka, J., Kong, E. H., Malivert, L., McIlwraith, M. J., Pape, T., West, S. C., and Zhang, X. (2014) Structure and mechanism of action of the BRCA2 breast cancer tumor suppressor. Nat. Struct. Mol. Biol. 21, 962–968.

(46) Xu, J., Zhao, L., Xu, Y., Zhao, W., Sung, P., and Wang, H. W. (2017) Cryo-EM structures of human RAD51 recombinase filaments during catalysis of DNA-strand exchange. Nat. Struct. Mol. Biol. 24, 40–46.

(47) Qiu, Y., Antony, E., Doganay, S., Ran Koh, H., Lohman, T. M., and Myong, S. (2013) Srs2 prevents Rad51 filament formation by repetitive motion on DNA. Nat. Commun. 4, 1–10.

(48) Zhao, W., Vaithiyalingam, S., San Filippo, J., Maranon, D. G., Jimenez-Sainz, J., Fontenay, G. V, Kwon, Y., Leung, S. G., Lu, L., Jensen, R. B., Chazin, W. J., Wiese, C., and Sung, P. (2015) Promotion of BRCA2-Dependent Homologous Recombination by DSS1 via RPA Targeting and DNA Mimicry. Mol. Cell 59, 176–187.

(49) Kaur, G., Ren, R., Hammel, M., Horton, J. R., Yang, J., Cao, Y., He, C., Lan, F., Lan, X., Blobel, G. A., Blumenthal, R. M., Zhang, X., and Cheng, X. (2023) Allosteric autoregulation of DNA binding via a DNA-mimicking protein domain: a biophysical study of ZNF410–DNA interaction using small angle X-ray scattering. Nucleic Acids Res. 51, 1674–1686.

(50) Berti, M., Teloni, F., Mijic, S., Ursich, S., Fuchs, J., Palumbieri, M. D., Krietsch, J., Schmid, J. A., Garcin, E. B., Gon, S., Modesti, M., Altmeyer, M., and Lopes, M. (2020) Sequential role of RAD51 paralog complexes in replication fork remodeling and restart. Nat. Commun. 11.

(51) Candelli, A., Holthausen, J. T., Depken, M., Brouwer, I., Franker, M. A. M., Marchetti, M., Heller, I., Bernard, S., Garcin, E. B., Modesti, M., Wyman, C., Wuite, G. J. L., and Peterman, E. J. G. (2014) Visualization and quantification of nascent RAD51 filament formation at single-monomer resolution. Proc. Natl. Acad. Sci. U. S. A. 111, 15090–15095.

