## supplement figures and table for "Structure of RADX and mechanism for regulation of RAD51 nucleofilaments"

### Slide 1
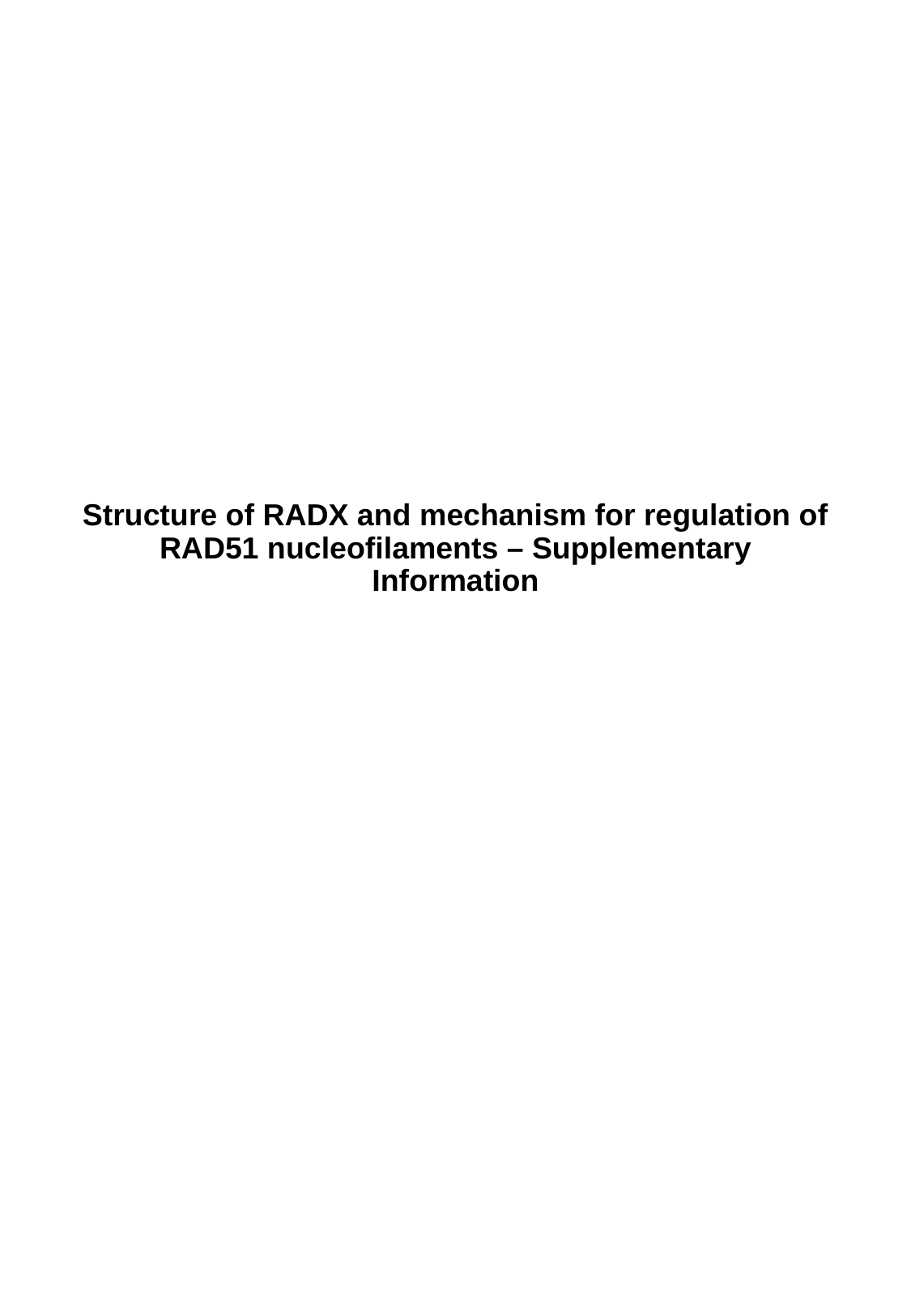

#### Structure of RADX and mechanism for regulation of RAD51 nucleofilaments – Supplementary Information

### Slide 2
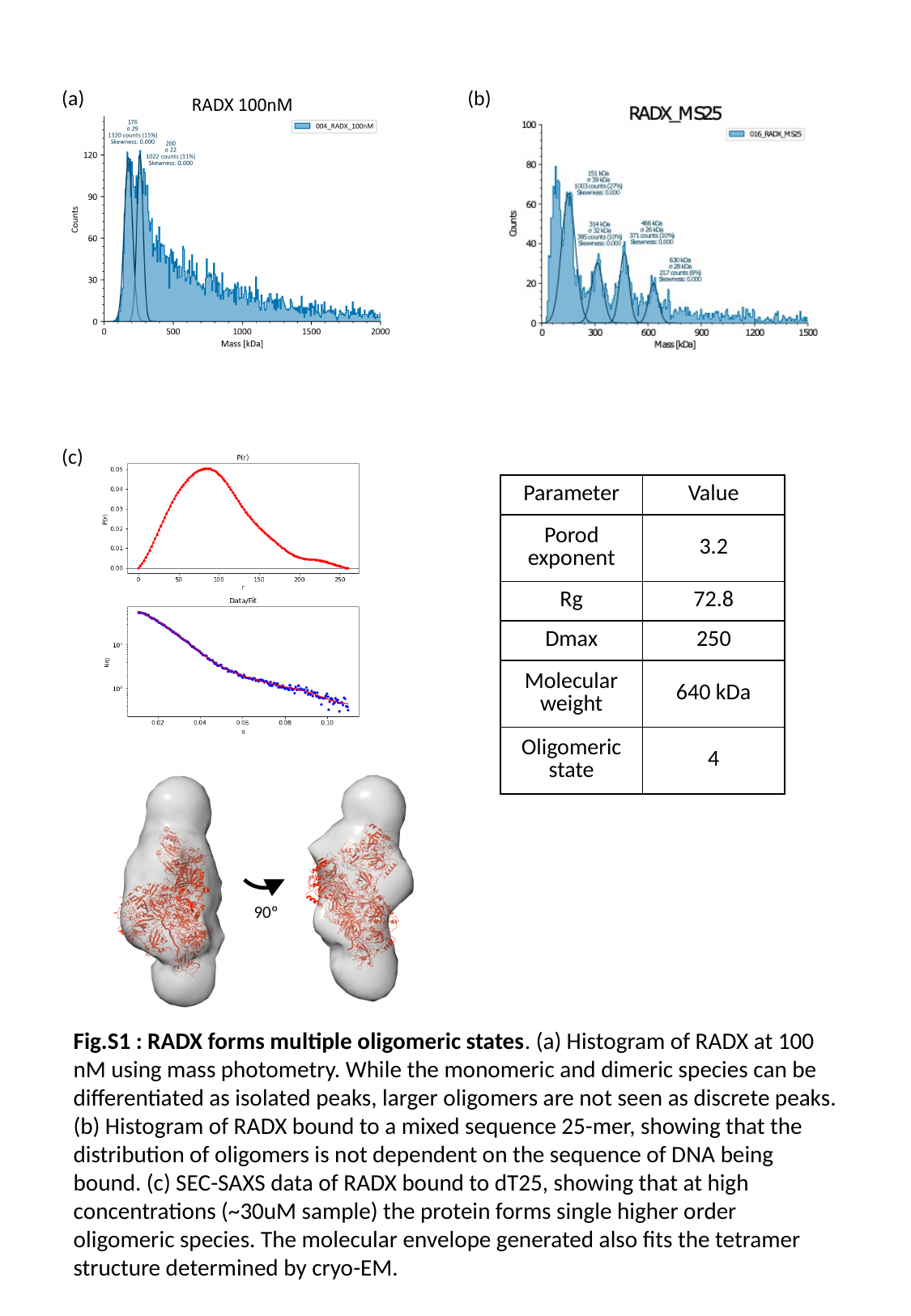

(b)
(a)
(c)
| Parameter | Value |
| --- | --- |
| Porod exponent | 3.2 |
| Rg | 72.8 |
| Dmax | 250 |
| Molecular weight | 640 kDa |
| Oligomeric state | 4 |
90º
Fig.S1 : RADX forms multiple oligomeric states. (a) Histogram of RADX at 100 nM using mass photometry. While the monomeric and dimeric species can be differentiated as isolated peaks, larger oligomers are not seen as discrete peaks. (b) Histogram of RADX bound to a mixed sequence 25-mer, showing that the distribution of oligomers is not dependent on the sequence of DNA being bound. (c) SEC-SAXS data of RADX bound to dT25, showing that at high concentrations (~30uM sample) the protein forms single higher order oligomeric species. The molecular envelope generated also fits the tetramer structure determined by cryo-EM.

### Slide 3
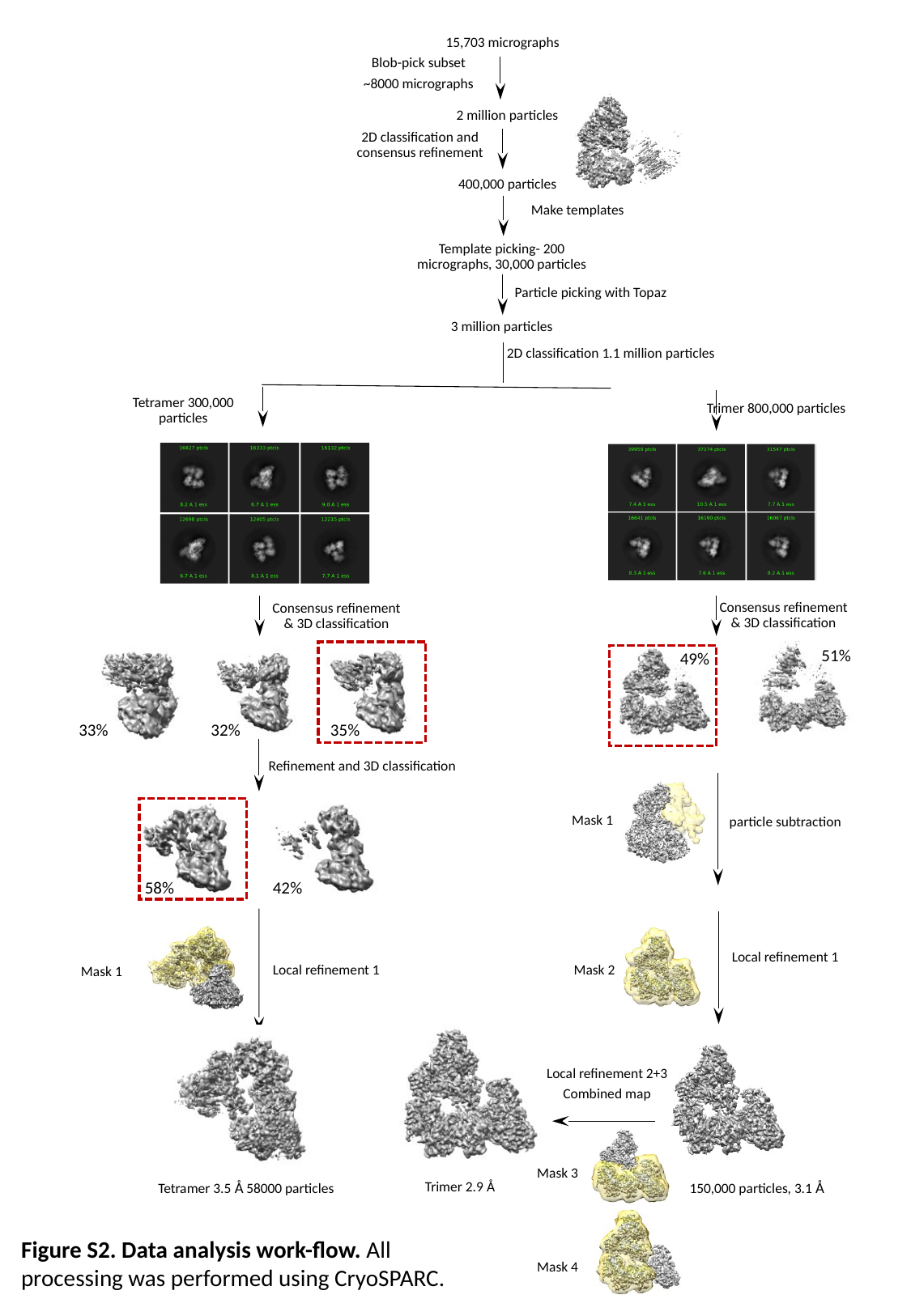

15,703 micrographs
Blob-pick subset
~8000 micrographs
2 million particles
2D classification and consensus refinement
400,000 particles
Make templates
Template picking- 200 micrographs, 30,000 particles
Particle picking with Topaz
3 million particles
2D classification 1.1 million particles
Trimer 800,000 particles
Tetramer 300,000 particles
Consensus refinement and 3D classification
Consensus refinement & 3D classification
Consensus refinement & 3D classification
51%
49%
33%
32%
35%
Refinement and 3D classification
Mask 1
particle subtraction
58%
42%
Local refinement 1
Local refinement 1
Mask 2
Mask 1
Local refinement 2+3
Combined map
Mask 3
 Trimer 2.9 Å
Tetramer 3.5 Å 58000 particles
150,000 particles, 3.1 Å
Figure S2. Data analysis work-flow. All processing was performed using CryoSPARC.
Mask 4

### Slide 4
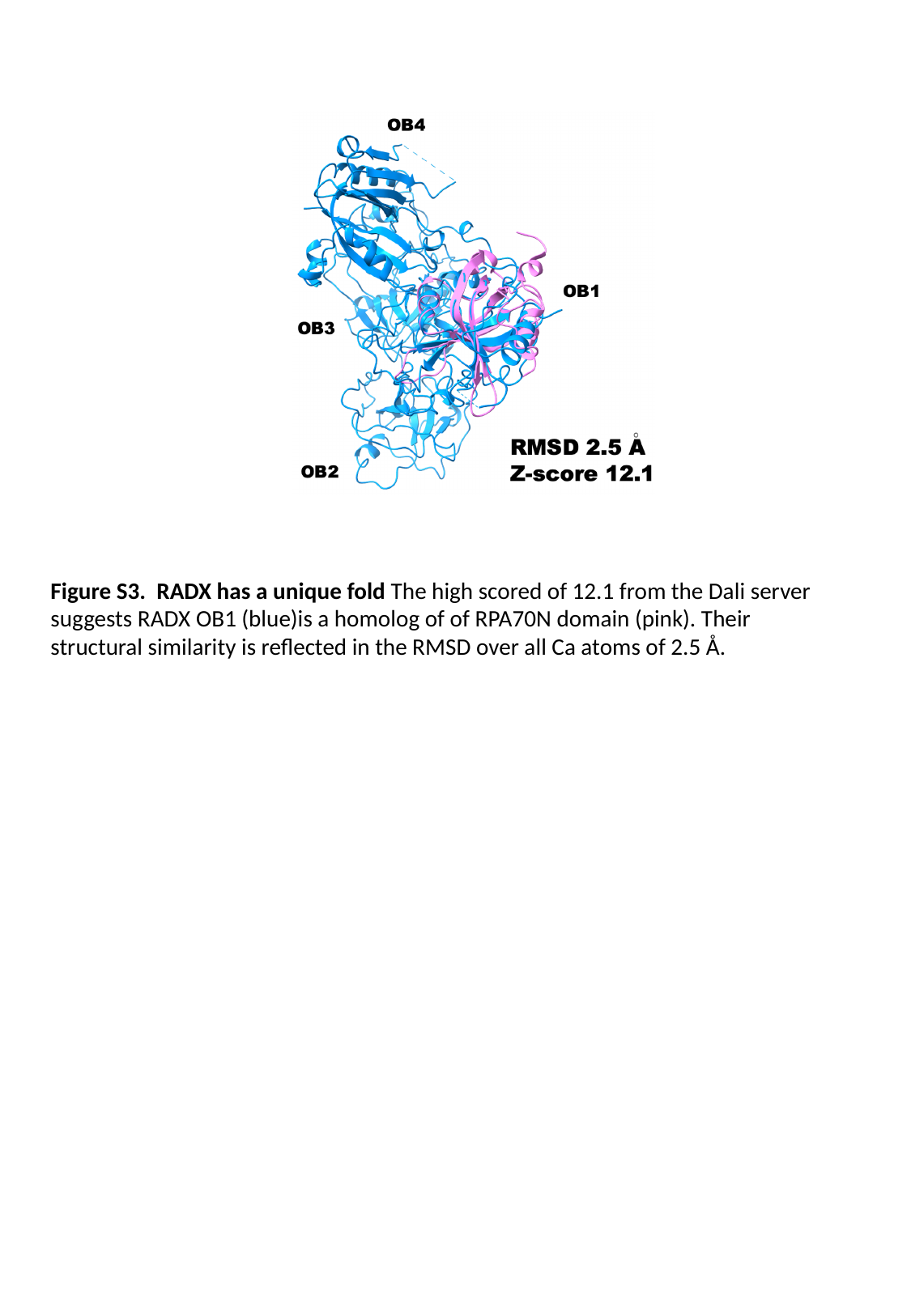

Figure S3. RADX has a unique fold The high scored of 12.1 from the Dali server suggests RADX OB1 (blue)is a homolog of of RPA70N domain (pink). Their structural similarity is reflected in the RMSD over all Ca atoms of 2.5 Å.

### Slide 5
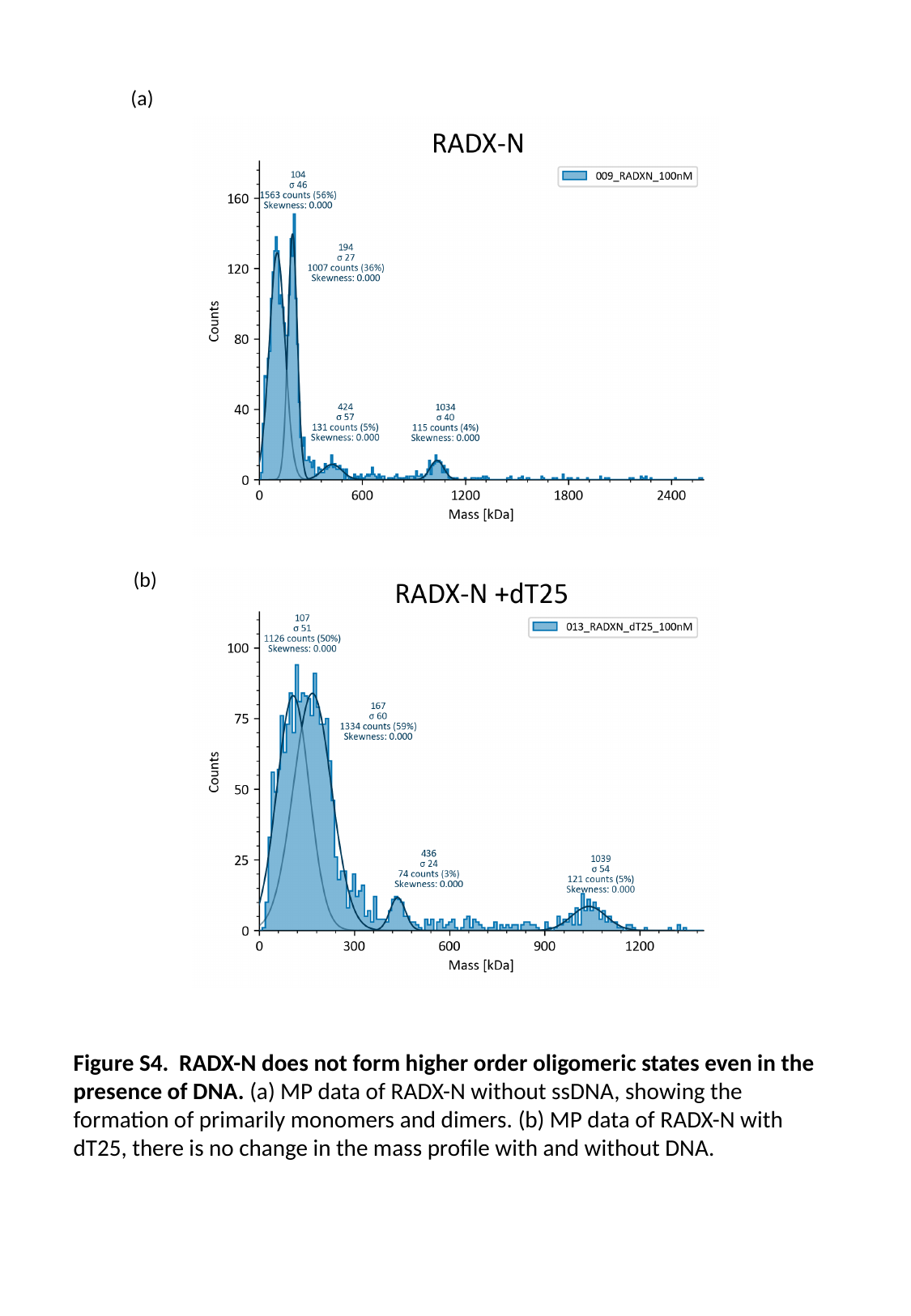

(a)
(b)
Figure S4. RADX-N does not form higher order oligomeric states even in the presence of DNA. (a) MP data of RADX-N without ssDNA, showing the formation of primarily monomers and dimers. (b) MP data of RADX-N with dT25, there is no change in the mass profile with and without DNA.

### Slide 6
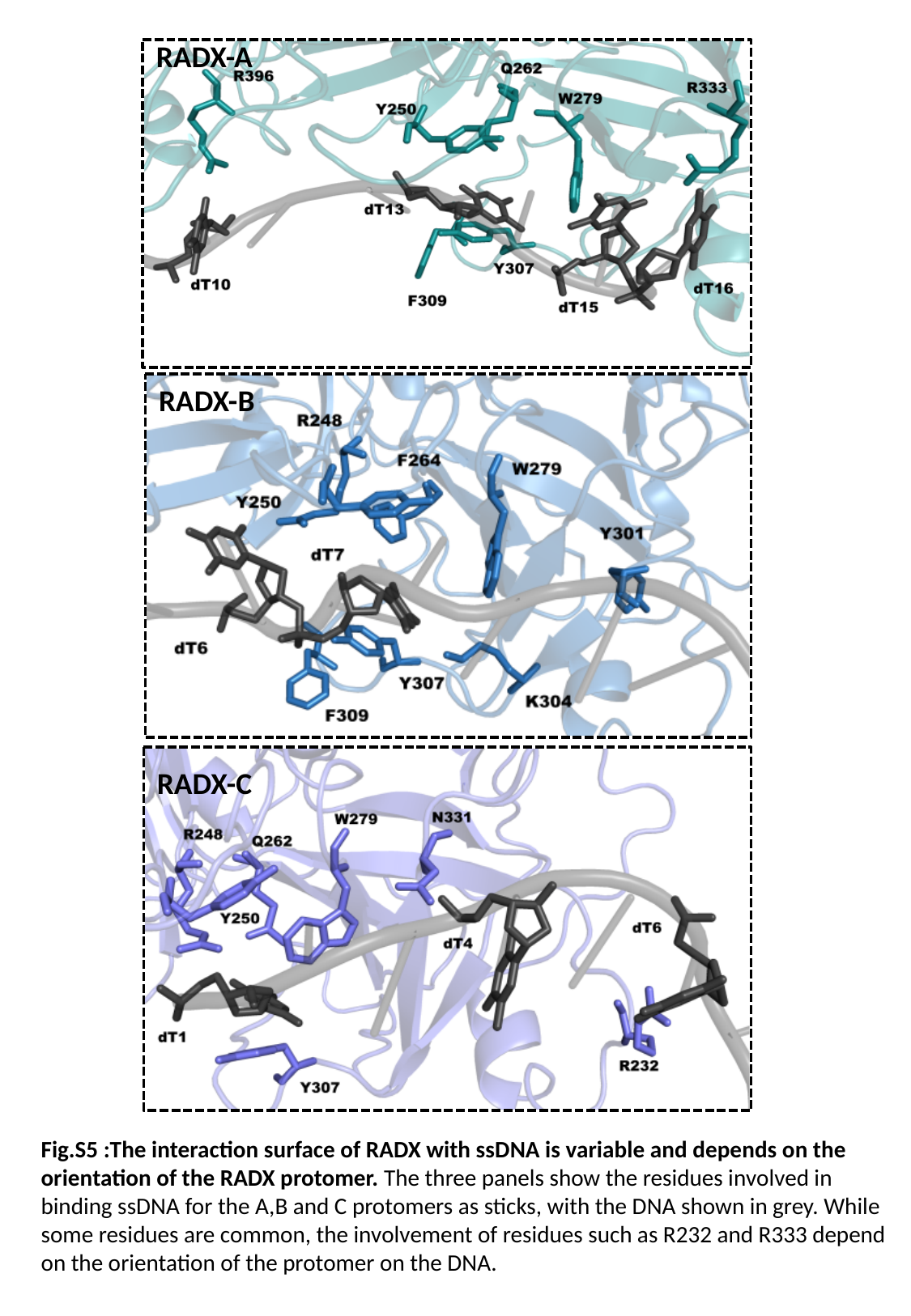

RADX-A
RADX-B
RADX-C
Fig.S5 :The interaction surface of RADX with ssDNA is variable and depends on the orientation of the RADX protomer. The three panels show the residues involved in binding ssDNA for the A,B and C protomers as sticks, with the DNA shown in grey. While some residues are common, the involvement of residues such as R232 and R333 depend on the orientation of the protomer on the DNA.

### Slide 7
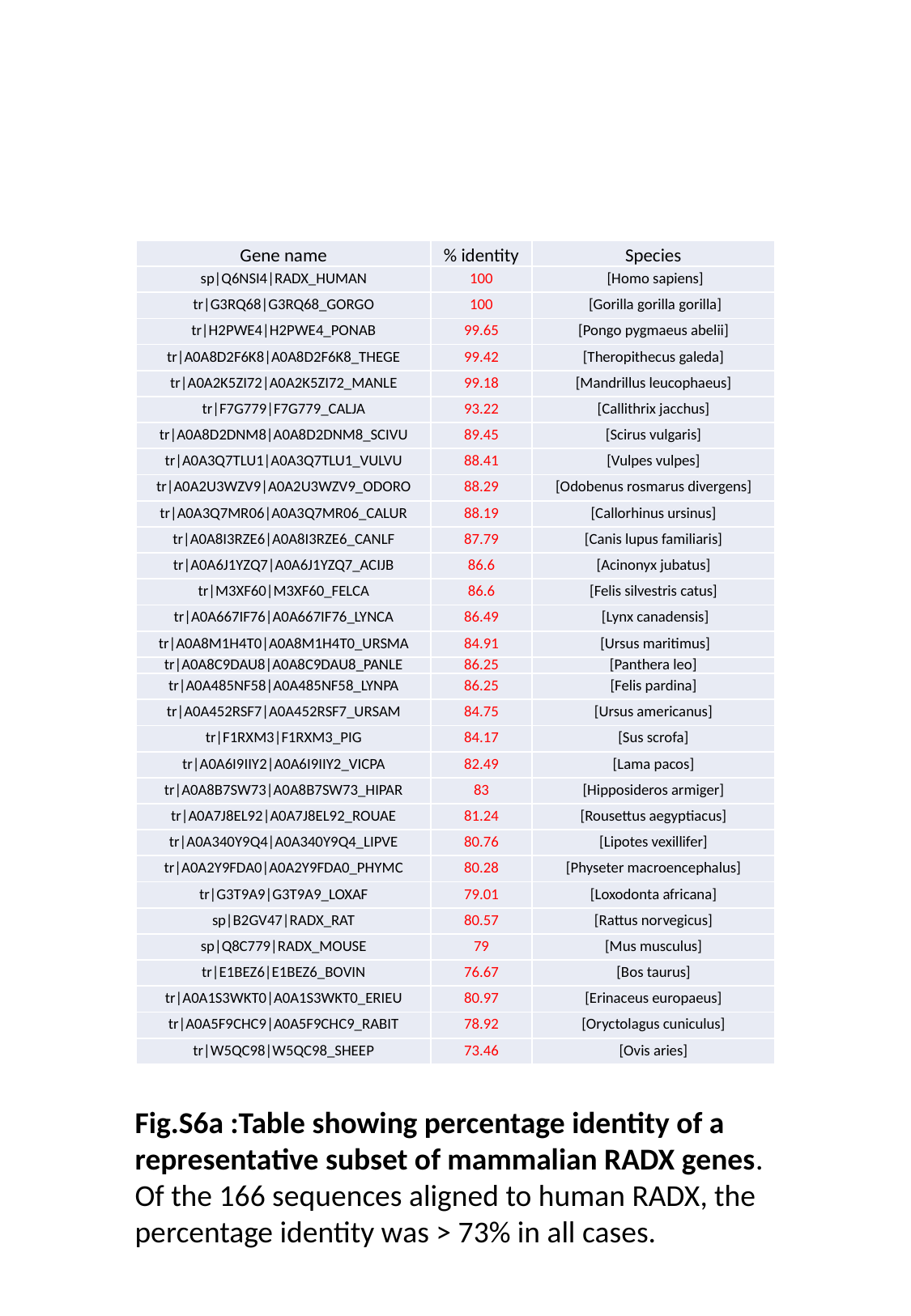

| Gene name | % identity | Species |
| --- | --- | --- |
| sp|Q6NSI4|RADX\_HUMAN | 100 | [Homo sapiens] |
| tr|G3RQ68|G3RQ68\_GORGO | 100 | [Gorilla gorilla gorilla] |
| tr|H2PWE4|H2PWE4\_PONAB | 99.65 | [Pongo pygmaeus abelii] |
| tr|A0A8D2F6K8|A0A8D2F6K8\_THEGE | 99.42 | [Theropithecus galeda] |
| tr|A0A2K5ZI72|A0A2K5ZI72\_MANLE | 99.18 | [Mandrillus leucophaeus] |
| tr|F7G779|F7G779\_CALJA | 93.22 | [Callithrix jacchus] |
| tr|A0A8D2DNM8|A0A8D2DNM8\_SCIVU | 89.45 | [Scirus vulgaris] |
| tr|A0A3Q7TLU1|A0A3Q7TLU1\_VULVU | 88.41 | [Vulpes vulpes] |
| tr|A0A2U3WZV9|A0A2U3WZV9\_ODORO | 88.29 | [Odobenus rosmarus divergens] |
| tr|A0A3Q7MR06|A0A3Q7MR06\_CALUR | 88.19 | [Callorhinus ursinus] |
| tr|A0A8I3RZE6|A0A8I3RZE6\_CANLF | 87.79 | [Canis lupus familiaris] |
| tr|A0A6J1YZQ7|A0A6J1YZQ7\_ACIJB | 86.6 | [Acinonyx jubatus] |
| tr|M3XF60|M3XF60\_FELCA | 86.6 | [Felis silvestris catus] |
| tr|A0A667IF76|A0A667IF76\_LYNCA | 86.49 | [Lynx canadensis] |
| tr|A0A8M1H4T0|A0A8M1H4T0\_URSMA | 84.91 | [Ursus maritimus] |
| tr|A0A8C9DAU8|A0A8C9DAU8\_PANLE | 86.25 | [Panthera leo] |
| tr|A0A485NF58|A0A485NF58\_LYNPA | 86.25 | [Felis pardina] |
| tr|A0A452RSF7|A0A452RSF7\_URSAM | 84.75 | [Ursus americanus] |
| tr|F1RXM3|F1RXM3\_PIG | 84.17 | [Sus scrofa] |
| tr|A0A6I9IIY2|A0A6I9IIY2\_VICPA | 82.49 | [Lama pacos] |
| tr|A0A8B7SW73|A0A8B7SW73\_HIPAR | 83 | [Hipposideros armiger] |
| tr|A0A7J8EL92|A0A7J8EL92\_ROUAE | 81.24 | [Rousettus aegyptiacus] |
| tr|A0A340Y9Q4|A0A340Y9Q4\_LIPVE | 80.76 | [Lipotes vexillifer] |
| tr|A0A2Y9FDA0|A0A2Y9FDA0\_PHYMC | 80.28 | [Physeter macroencephalus] |
| tr|G3T9A9|G3T9A9\_LOXAF | 79.01 | [Loxodonta africana] |
| sp|B2GV47|RADX\_RAT | 80.57 | [Rattus norvegicus] |
| sp|Q8C779|RADX\_MOUSE | 79 | [Mus musculus] |
| tr|E1BEZ6|E1BEZ6\_BOVIN | 76.67 | [Bos taurus] |
| tr|A0A1S3WKT0|A0A1S3WKT0\_ERIEU | 80.97 | [Erinaceus europaeus] |
| tr|A0A5F9CHC9|A0A5F9CHC9\_RABIT | 78.92 | [Oryctolagus cuniculus] |
| tr|W5QC98|W5QC98\_SHEEP | 73.46 | [Ovis aries] |
Fig.S6a :Table showing percentage identity of a representative subset of mammalian RADX genes. Of the 166 sequences aligned to human RADX, the percentage identity was > 73% in all cases.

### Slide 8
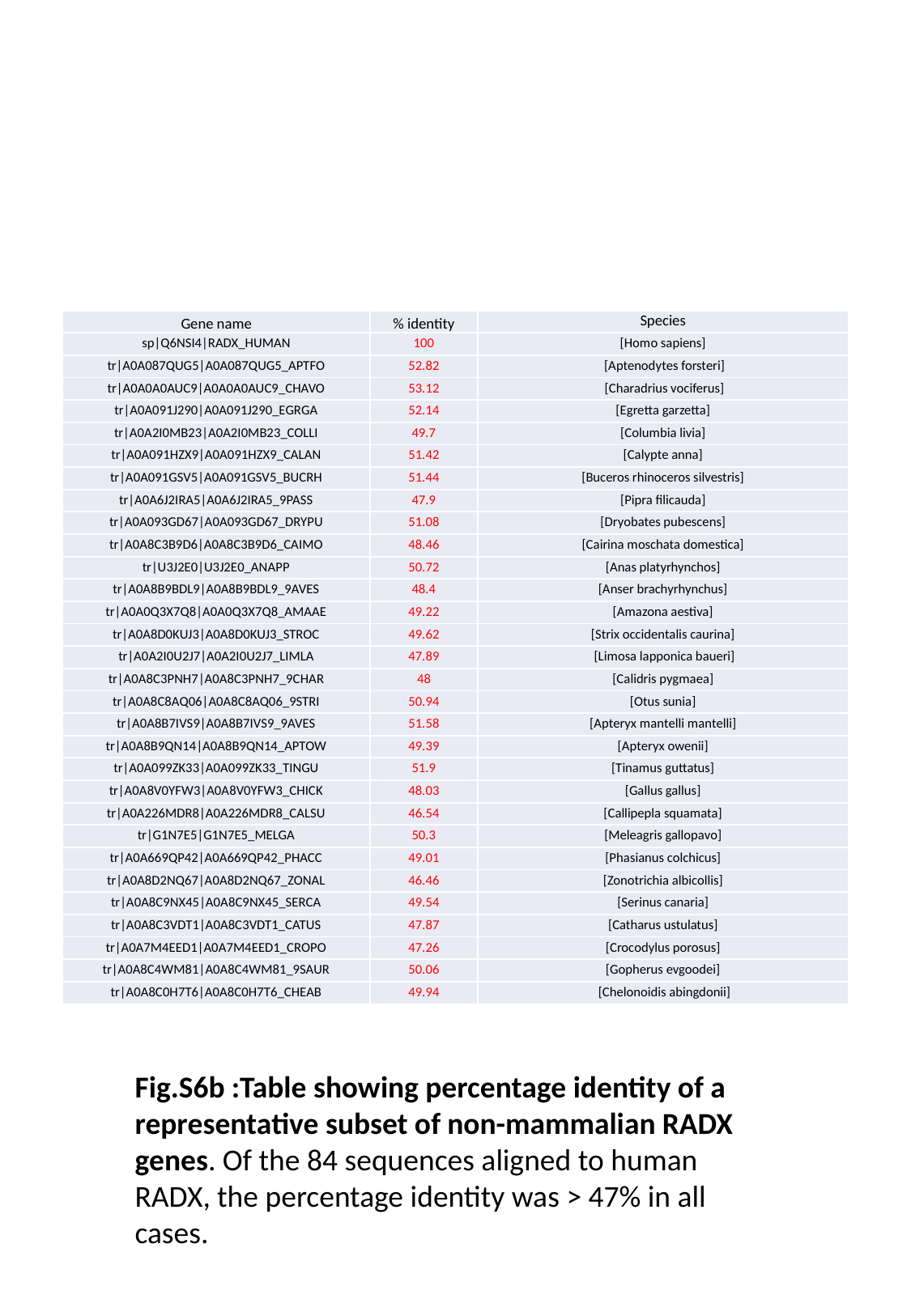

| Gene name | % identity | Species |
| --- | --- | --- |
| sp|Q6NSI4|RADX\_HUMAN | 100 | [Homo sapiens] |
| tr|A0A087QUG5|A0A087QUG5\_APTFO | 52.82 | [Aptenodytes forsteri] |
| tr|A0A0A0AUC9|A0A0A0AUC9\_CHAVO | 53.12 | [Charadrius vociferus] |
| tr|A0A091J290|A0A091J290\_EGRGA | 52.14 | [Egretta garzetta] |
| tr|A0A2I0MB23|A0A2I0MB23\_COLLI | 49.7 | [Columbia livia] |
| tr|A0A091HZX9|A0A091HZX9\_CALAN | 51.42 | [Calypte anna] |
| tr|A0A091GSV5|A0A091GSV5\_BUCRH | 51.44 | [Buceros rhinoceros silvestris] |
| tr|A0A6J2IRA5|A0A6J2IRA5\_9PASS | 47.9 | [Pipra filicauda] |
| tr|A0A093GD67|A0A093GD67\_DRYPU | 51.08 | [Dryobates pubescens] |
| tr|A0A8C3B9D6|A0A8C3B9D6\_CAIMO | 48.46 | [Cairina moschata domestica] |
| tr|U3J2E0|U3J2E0\_ANAPP | 50.72 | [Anas platyrhynchos] |
| tr|A0A8B9BDL9|A0A8B9BDL9\_9AVES | 48.4 | [Anser brachyrhynchus] |
| tr|A0A0Q3X7Q8|A0A0Q3X7Q8\_AMAAE | 49.22 | [Amazona aestiva] |
| tr|A0A8D0KUJ3|A0A8D0KUJ3\_STROC | 49.62 | [Strix occidentalis caurina] |
| tr|A0A2I0U2J7|A0A2I0U2J7\_LIMLA | 47.89 | [Limosa lapponica baueri] |
| tr|A0A8C3PNH7|A0A8C3PNH7\_9CHAR | 48 | [Calidris pygmaea] |
| tr|A0A8C8AQ06|A0A8C8AQ06\_9STRI | 50.94 | [Otus sunia] |
| tr|A0A8B7IVS9|A0A8B7IVS9\_9AVES | 51.58 | [Apteryx mantelli mantelli] |
| tr|A0A8B9QN14|A0A8B9QN14\_APTOW | 49.39 | [Apteryx owenii] |
| tr|A0A099ZK33|A0A099ZK33\_TINGU | 51.9 | [Tinamus guttatus] |
| tr|A0A8V0YFW3|A0A8V0YFW3\_CHICK | 48.03 | [Gallus gallus] |
| tr|A0A226MDR8|A0A226MDR8\_CALSU | 46.54 | [Callipepla squamata] |
| tr|G1N7E5|G1N7E5\_MELGA | 50.3 | [Meleagris gallopavo] |
| tr|A0A669QP42|A0A669QP42\_PHACC | 49.01 | [Phasianus colchicus] |
| tr|A0A8D2NQ67|A0A8D2NQ67\_ZONAL | 46.46 | [Zonotrichia albicollis] |
| tr|A0A8C9NX45|A0A8C9NX45\_SERCA | 49.54 | [Serinus canaria] |
| tr|A0A8C3VDT1|A0A8C3VDT1\_CATUS | 47.87 | [Catharus ustulatus] |
| tr|A0A7M4EED1|A0A7M4EED1\_CROPO | 47.26 | [Crocodylus porosus] |
| tr|A0A8C4WM81|A0A8C4WM81\_9SAUR | 50.06 | [Gopherus evgoodei] |
| tr|A0A8C0H7T6|A0A8C0H7T6\_CHEAB | 49.94 | [Chelonoidis abingdonii] |
Fig.S6b :Table showing percentage identity of a representative subset of non-mammalian RADX genes. Of the 84 sequences aligned to human RADX, the percentage identity was > 47% in all cases.

### Slide 9
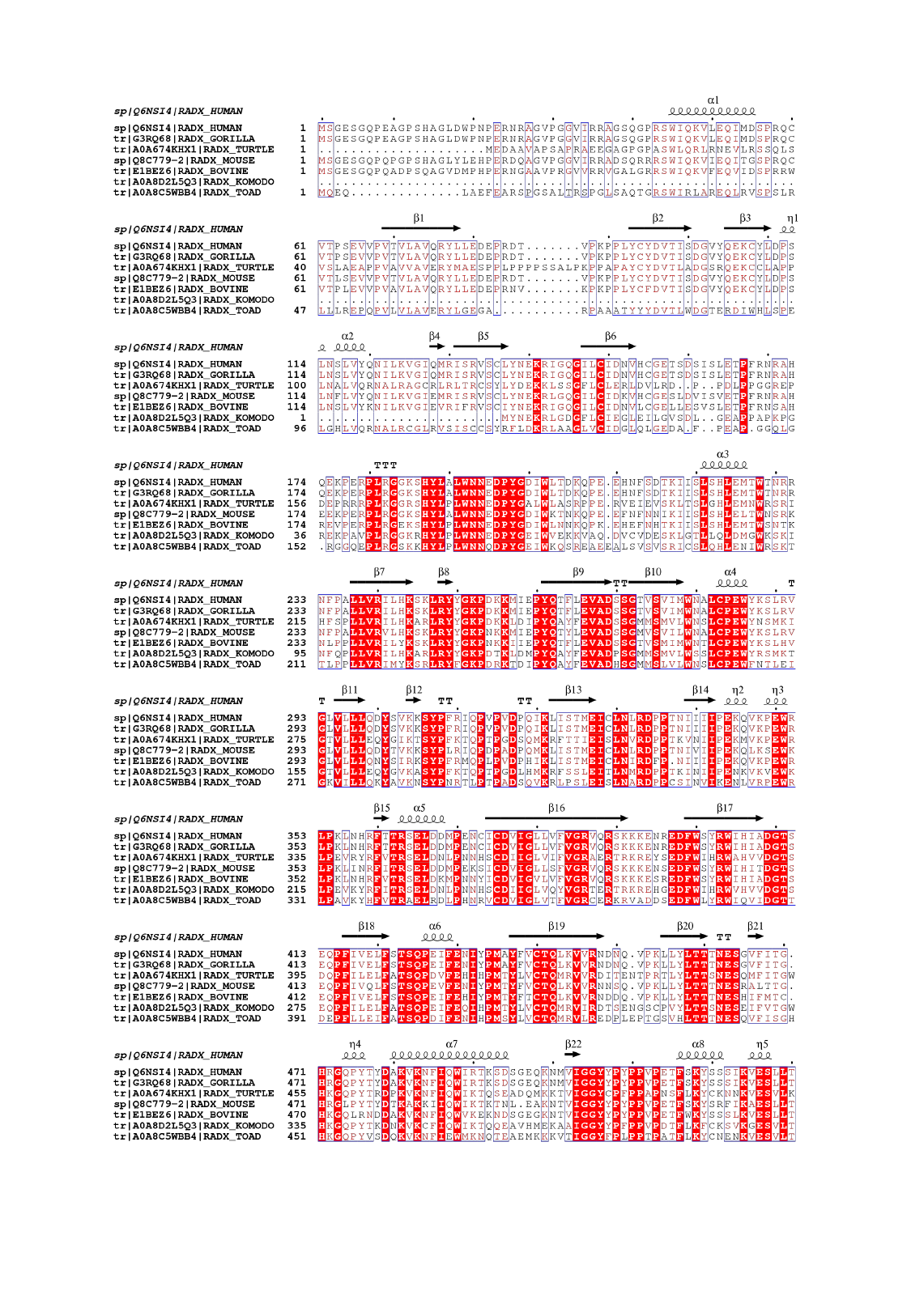

### Slide 10
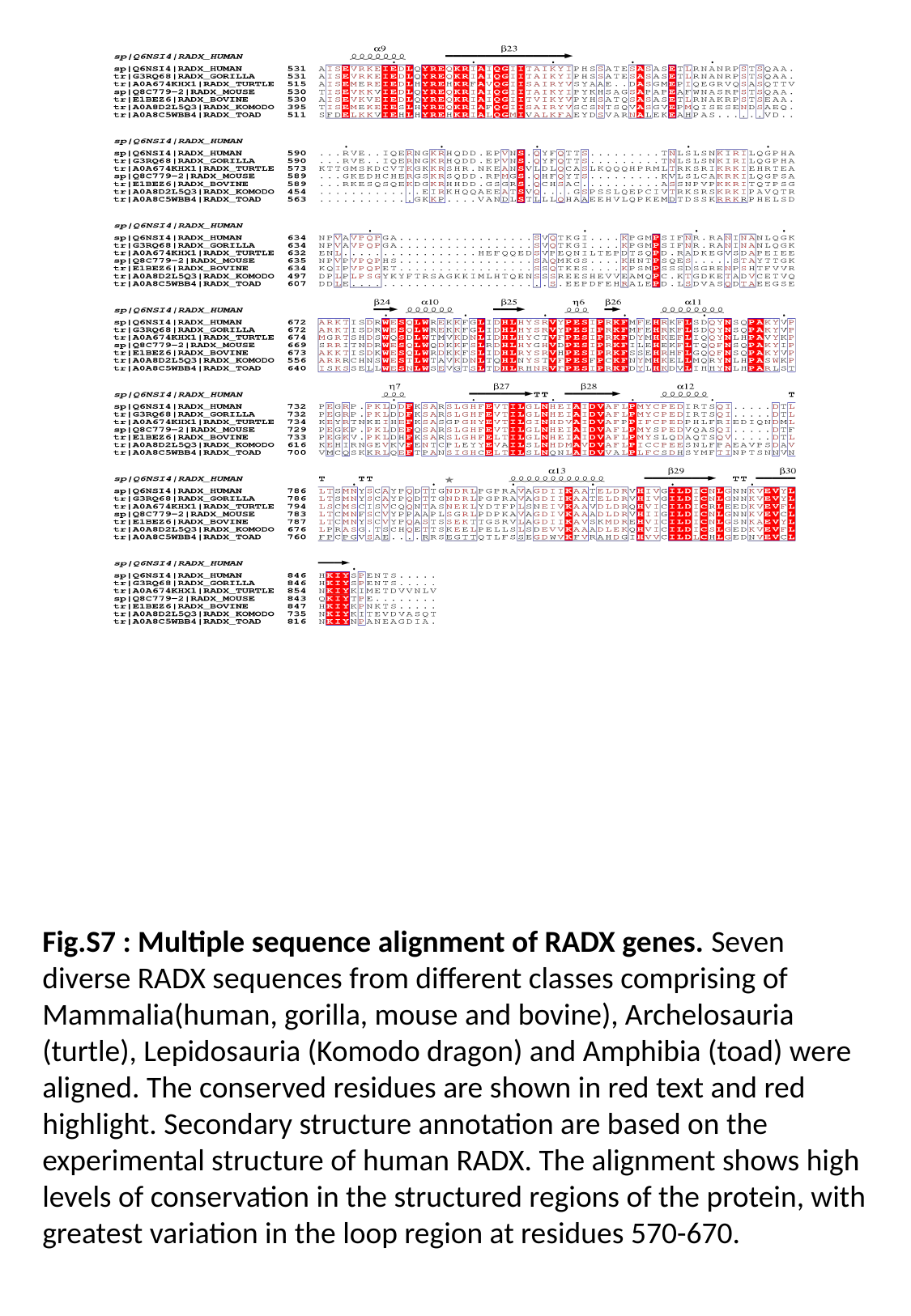

Fig.S7 : Multiple sequence alignment of RADX genes. Seven diverse RADX sequences from different classes comprising of Mammalia(human, gorilla, mouse and bovine), Archelosauria (turtle), Lepidosauria (Komodo dragon) and Amphibia (toad) were aligned. The conserved residues are shown in red text and red highlight. Secondary structure annotation are based on the experimental structure of human RADX. The alignment shows high levels of conservation in the structured regions of the protein, with greatest variation in the loop region at residues 570-670.

### Slide 11
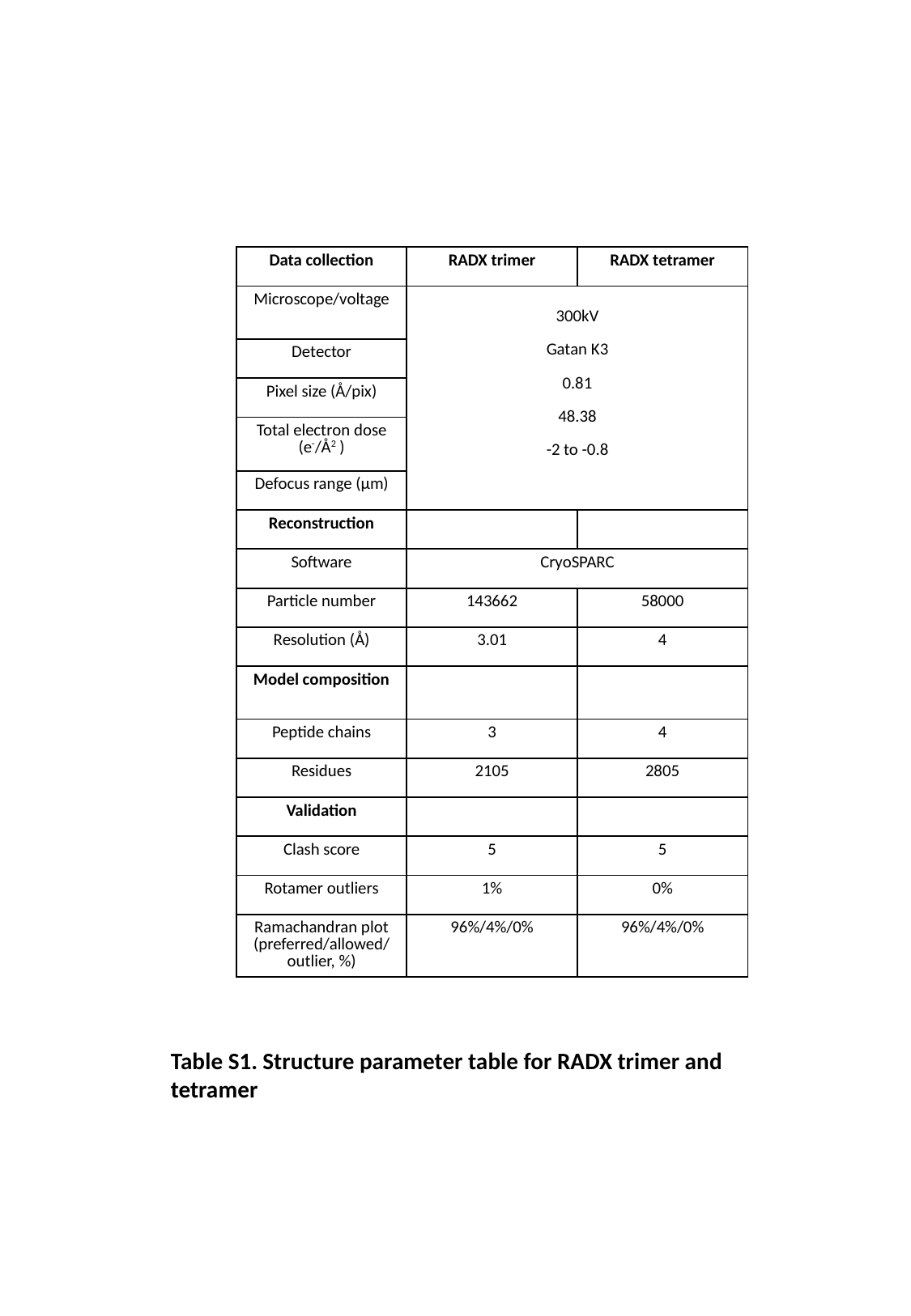

| Data collection | RADX trimer | RADX tetramer |
| --- | --- | --- |
| Microscope/voltage | 300kV Gatan K3 0.81 48.38 -2 to -0.8 | 300kV |
| Detector | Gatan K3 | Gatan K3 |
| Pixel size (Å/pix) | 0.81 | 0.81 |
| Total electron dose (e^-^/Å^2^ ) | 48.38 | 48.38 |
| Defocus range (µm) | -2 to -0.8 | -2 to -0.8 |
| Reconstruction | | |
| Software | CryoSPARC | CryoSPARC |
| Particle number | 143662 | 58000 |
| Resolution (Å) | 3.01 | 4 |
| Model composition | | |
| Peptide chains | 3 | 4 |
| Residues | 2105 | 2805 |
| Validation | | |
| Clash score | 5 | 5 |
| Rotamer outliers | 1% | 0% |
| Ramachandran plot (preferred/allowed/outlier, %) | 96%/4%/0% | 96%/4%/0% |
Table S1. Structure parameter table for RADX trimer and tetramer
